# Tumor-derived biomarkers beyond antigen expression enhance efficacy of CD276/B7H3 antibody-drug conjugate in metastatic prostate cancer

**DOI:** 10.1101/2022.04.19.488784

**Authors:** Supreet Agarwal, Lei Fang, Kerry McGowen, JuanJuan Yin, Joel Bowman, Anson T. Ku, Aian Neil Alilin, Eva Corey, Martine Roudier, Lawrence True, Ruthy Dumpit, Ilsa Coleman, John Lee, Peter S. Nelson, Brian J. Capaldo, Aida Mariani, Clare Hoover, Ilya S. Senatorov, Michael Beshiri, Adam G. Sowalsky, Elaine M Hurt, Kathleen Kelly

## Abstract

Antibody-drug conjugates (ADCs) are promising targeted cancer therapy; however, patient selection based solely on target antigen expression without consideration for cytotoxic payload vulnerabilities has plateaued clinical benefits. Biomarkers to capture patients who might benefit from specific ADCs have not been systematically determined for any cancer. We present a comprehensive therapeutic and biomarker analysis of a B7H3-ADC with pyrrolobenzodiazepine(PBD) payload in 26 treatment-resistant, metastatic prostate cancer(mPC) models. B7H3 is a tumor-specific surface protein widely expressed in mPC, and PBD is a DNA cross-linking agent. B7H3 expression was necessary but not sufficient for B7H3-PBD-ADC responsiveness. RB1 deficiency and/or replication stress, characteristics of poor prognosis, conferred sensitivity and were associated with complete tumor regression in both neuroendocrine (NEPC) and androgen receptor positive(ARPC) prostate cancer models, even with low B7H3 levels. Non-ARPC models, which are currently lacking efficacious treatment, demonstrated the highest replication stress and were most sensitive to treatment. In RB1 wild-type ARPC tumors, SLFN11 expression or select DNA repair mutations in SLFN11 non-expressors governed response. Significantly, wild-type TP53 predicted non-responsiveness (7/8 models). Overall, biomarker-focused selection of models led to high efficacy of in vivo treatment. These data enable a paradigm shift to biomarker-driven trial designs for maximizing clinical benefit of ADC therapies.

## INTRODUCTION

Metastatic prostate cancer (mPC) remains a lethal disease accounting for more than 30,000 deaths annually in the United States (1). The use of androgen deprivation therapy (ADT) and androgen receptor (AR) signaling inhibitors (ARSI) have significantly prolonged the survival of patients with mPC in both pre- and post-chemotherapy settings. Unfortunately, durable complete responses are uncommon and mortality rates approach 100% with the development of castration resistant prostate cancer (mCRPC). Continuously evolving acquired resistance mechanisms include frequent AR mutations and structural genomic alterations that drive an AR-active adenocarcinoma phenotype (ARPC). Less common, but increasing in frequency, are resistance mechanisms that bypass an AR requirement through lineage plasticity with the emergence of phenotypes spanning neuroendocrine phenotypes (NEPC) and various other histologies (2). Across the landscape of genomic alterations in mCRPC, *RB1* alteration is the only genomic factor strongly associated with poor survival (3), highlighting the need for potential therapeutic strategies targeting RB1 deficient tumors. Since the vast majority of mPC phenotypes eventually resist all currently approved therapeutics, new treatment strategies are essential.

A promising approach for developing effective and less toxic therapies for mPC involves selectively targeting tumor cells via tumor-specific cell surface proteins and cognate antigens. Exploiting prostate specific membrane antigen (PSMA) to deliver high dose radiation to tumor cells overexpressing PSMA, PSMA-Lu177, has recently gained FDA approval for the treatment of mPC (4). PSMA is expressed by the majority, though not all AR active PCs, but emerging treatment-resistant phenotypes such as AR-negative and small cell neuroendocrine PC (SCNPC) generally do not express PSMA, prompting a search for alternate targets. CD276/B7H3 is a type I transmembrane protein overexpressed in several solid tumors and often correlated with poor survival and higher tumor grade (5, 6). B7H3 is overexpressed in prostate cancer compared to benign prostatic hyperplasia, and high B7H3 expression is positively correlated with adenocarcinoma aggressiveness, observed as overexpression in metastatic and castrate resistant disease (7–9). Further, B7H3 expression is not detected in human normal pancreas, lung, liver, kidney, colon, and heart (10). The preferential overexpression of B7H3 protein on the surface of cancer cells and the minimal expression on normal tissues makes it an ideal target for antibody-based therapeutics (11, 12), and targeting B7H3 is being widely pursued as more than 30 clinical trials are currently registered on clinicaltrials.gov.

One common strategy utilizing cell surface targets such as PSMA, B7H3, PSCA, TROP2, STEAP1, and CEACAM5 include the development of antibody drug conjugates (ADCs) (13, 14). ADCs combine the high target specificity of a monoclonal antibody with a cytotoxic agent for targeted killing of tumor cells. Several classes of cytotoxic drugs have been utilized in ADC designs, though most are either potent microtubule poisons or inducers of DNA damage (15). ADCs are rapidly internalized, releasing the antibody-linked payload to induce cell death. Several ADCs have been evaluated preclinically; however, only a few have been approved for clinical use due to either lack of efficacy or unacceptable toxicity, highlighting the need for strategies to define and use criteria for patient selection (16, 17). Generally, ADC based trials have primarily focused on target antigen expression as a patient selection strategy, but absolute levels of target antigen have not been sufficient to predict response, suggesting that multiple factors be considered, including underlying mechanisms of vulnerability toward the cytotoxic payload (17). In fact, FDA approved HER2 targeted ADC (Enhertu), has shown efficacy in metastatic HER2-low breast cancer subtype (indication revised in August 2022) clearly suggesting a need for biomarkers other than the target antigen. Thus, a composite set of biomarkers including target protein expression are required to maximize clinical benefits of ADCs.

Pyrrolobenzodiazepines (PBD) are DNA minor-groove crosslinking agents that have been used as payloads for several clinical grade ADCs (18). PBD dimers bind in a sequence specific manner to form inter-strand crosslinks (ICLs) leading to double strand DNA breaks due to replication fork arrest (18). Determining the utility of PBD dimers for the treatment of cancers exhibiting replication stress and consequently increased potential vulnerability is a question of interest (19). Further, how tumor heterogeneity influences payload sensitivity and therapeutic efficacy has not been broadly investigated in preclinical cohorts of defined tumor types.

In this study, we evaluated a therapeutic strategy targeting B7H3 for the treatment of mPC. We profiled a spectrum of mPC tumors to assess the heterogeneity of B7H3 expression with respect to metastatic site and tumor phenotype. We tested a humanized B7H3-ADC armed with PBD payload across a range of molecularly characterized, clinically relevant mPC PDX and organoid models that reflect the diversity of human mPCs. Although we anticipated that DNA-double strand break repair defects and levels of B7H3 expression would drive B7H3-PBD-ADC responses, we observed high efficacy in 1) select B7H3-low expressing models with no apparent mutations in DNA repair pathway genes, and 2) no response in a group of B7H3^+^ adenocarcinomas. By integrating genomic and transcriptomic characteristics with B7H3-PBD-ADC response data, we uncovered additional biomarkers that represent vulnerabilities derived from more than one sensitivity or resistance pathway. These analyses demonstrate how a diverse cohort of mPCs distribute into distinct biomarker classes that reflect ADC mechanisms of action. Collectively the results have the potential to inform patient selection for prospective trials and contribute to the interpretation of patient response and resistance outcomes.

## RESULTS

### B7H3 is expressed across a range of mPC phenotypes and diverse metastatic sites

To assess the potential clinical utility of targeting B7H3 as a treatment strategy for advanced prostate cancer, we evaluated the transcript abundance of B7H3 and other previously studied cell surface targets (PSMA, PSCA, TROP2, STEAP1, and CEACAM5) in 185 tumors from 98 treatment refractory prostate cancer metastasis (mPC) patients, and across a panel of 26 mPC PDX and organoid models representing the genomic and phenotypic heterogeneity of patient tumors. The 26 preclinical models tested in this study comprise tumors of adenocarcinoma (ARPC) phenotype with AR signaling (intact: n= 13, experimentally castrate resistant (CR): n=6) as well as AR^NEG/VERY^ ^LOW^ non-neuroendocrine prostate cancer (n=2), denoted as DNPC, (2) and small cell neuroendocrine prostate cancer (n=5), denoted at SCNPC. The latter two groups were collectively categorized as non-ARPC (**Figure S1A)**. The models included were primarily from the LuCaP PDX series (20) and also included 2 NCI mPC patient biopsy-derived organoids (PDOs) of the ARPC phenotype (21). We also categorized each patient tumor and mPC model into phenotypic categories based on gene expression signatures reflecting AR signaling and neuroendocrine (NE) pathway activity. We first quantified *CD276(B7H3)* transcript abundance in patient samples (185 tumors from 98 mPC patients) and the above described 26 mPC models using RNAseq measurements. Overall, the vast majority of samples expressed *CD276* transcripts, and there was limited variation within or between mPC phenotypes compared to other targets (**Figure 1, A-B and Figure S1B**). *CD276* was also the most consistently expressed target across different mCRPC phenotypes. In contrast, other markers such as *FOLH1(PSMA)* expression varied substantially both within a phenotype and between phenotypes *(P=*1×10^-9^ for the mean Log_2_ FKPM values between ARPC vs. NEPC) **(Figure 1B:** *FOLH1* (blue), *CD276*(green)). We also evaluated the intra-individual heterogeneity of *CD276* transcript levels in multiple tumors acquired from the same patient. With few exceptions, there was a tight distribution of *CD276* expression within individuals (**Figure S1C**).

**FIGURE 1.**
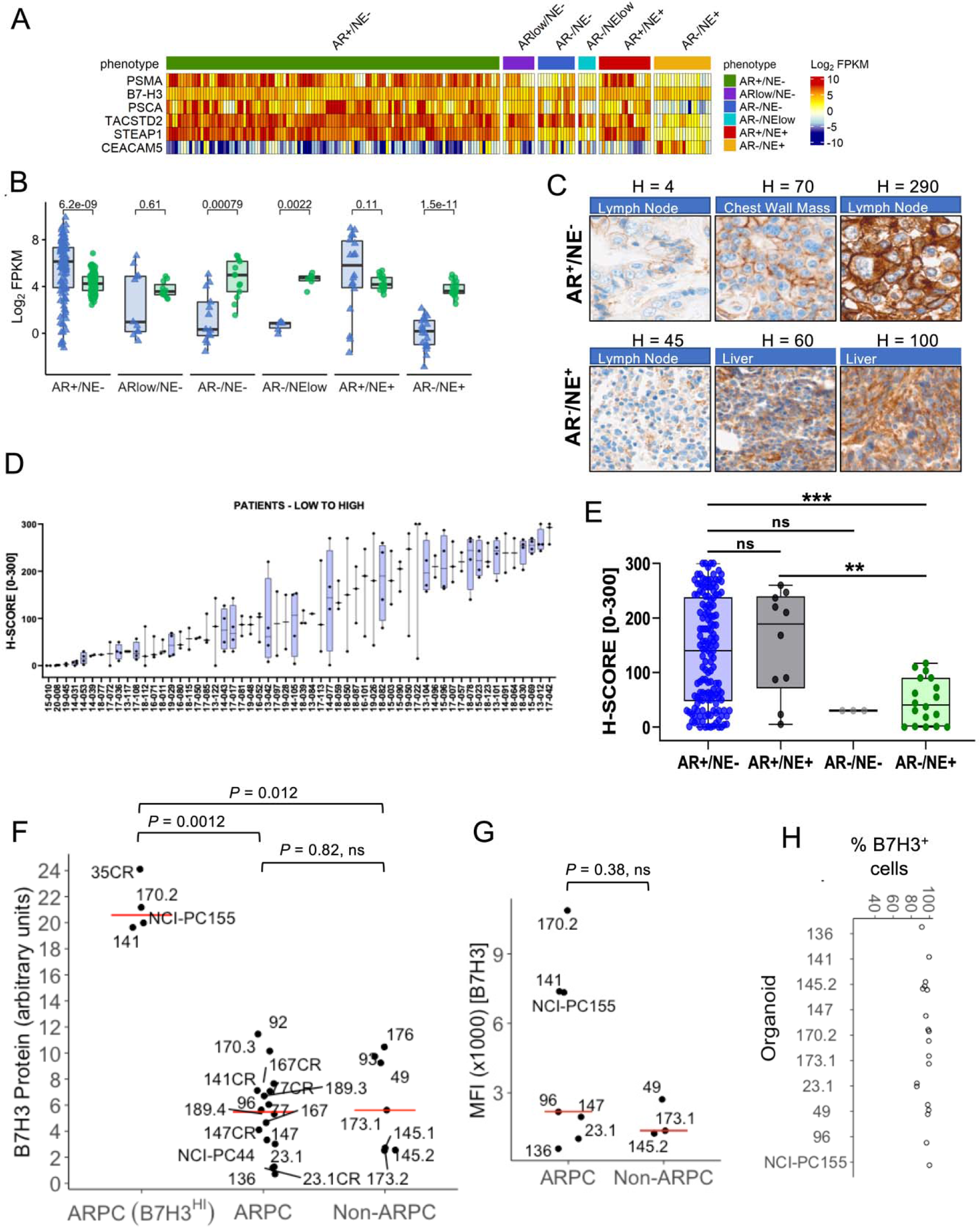
CD276/B7H3 expression in metastatic prostate cancer patient samples and mPC PDX/organoid models. **(A)** *CD276/B7H3, FOLH1/PSMA, PSCA, TACSTD2/TROP2, STEAP1*, and *CEACAM5* transcript abundance determined by RNA sequencing analysis of 185 metastatic prostate tumors from 98 patients. Transcript levels are shown as Log2 FPKM. **(B)** Comparisons of *CD276/B7H3* (green dots) and *FOLH1/PSMA* (blue dots) expression by phenotypes of metastatic tumors. Groups were compared using two-sided Wilcoxon rank tests with Benjamini-Hochberg multiple-testing correction. **(C)** Immunohistochemical assessments of B7H3 protein expression. Representative staining of tumors with low, medium, and high B7H3 expression in AR+/NE- and AR-/NE+ phenotypes. **(D)** Distribution of B7H3 protein expression in 181 metastatic tumors within and between 58 patients. **(E)** Distribution of B7H3 protein expression in metastatic prostate cancers categorized by phenotype. **(F)** Western blot quantification of B7H3 protein expression in PDX tissue samples and two PDOs (NCI-PC44, NCI-PC155) by Simple Western. ARPC samples with high B7H3 expression are categorized separately in the B7H3^HI^ group. Y-axis represents CD276/B7H3 protein quantification scaled by a factor of 10. ns = not significant. **(G-H)** Flow cytometry analysis for B7H3 cell-surface expression from organoids dissociated into single cells. **(G)** Median Fluorescence Intensity (MFI) and **(H)** % positive cells are shown for nine analyzed models.

We next evaluated B7H3 protein expression across a cohort of PC metastases using a tissue microarray comprised of 181 tumors from 58 patients (range of 1 to 4 tumors per patient). Three tumors were not analyzed due to insufficient tumor content, leaving 178 evaluable tumors.

Overall, B7H3 protein exhibited more variation compared to transcript expression: of 178 tumors evaluated, 149 expressed B7H3 (H-score > 20) and 29 lacked expression (**Figure 1, C and D**). B7H3 was detected across diverse metastatic sites with bone metastases exhibiting the highest levels (**Figure S1D**). Tumors categorized as AR+/NE-ARPC generally expressed higher B7H3 levels compared to other phenotypes, but a subset of AR-/NE+ SCNPC and AR-/NE-tumors also expressed B7H3 (**Figure 1E**). Collectively, these results indicate that B7H3 may represent a target for antigen-directed therapeutics across a range of clinical mPC phenotypes.

Additionally, we used a quantitative immunoblot technique to determine the relative amount of total B7H3 protein expressed in mPC preclinical models. Like patient samples, we observed wide variation (>30-fold) in B7H3 protein levels (**Figure 1F** and **Figure S1E**). ARPC models demonstrated a range of expression clustering as a high group (B7H3^Hi^) and an intermediate to low group, the latter of which overlapped in median level with the non-ARPC group (**Figure 1F**). There was no apparent common genotypic or phenotypic feature in B7H3^HI^ ARPC group. Consistent with patient data, B7H3 mRNA levels were not strongly correlated with B7H3 protein levels in models of either phenotype (**Figure S1F**) emphasizing the minimal utility in transcriptional based assays for quantitative analyses (12). Importantly, in FACS analysis, despite variability in the median fluorescence intensity across the models tested (n=9), EpCam+ tumor cells homogenously expressed B7H3 (80-100% cells) at the cell surface which makes B7H3 an ideal target for ADC based therapy (**Figure 1, G and H**, and **Figure S2A**). B7H3 cell surface level was well correlated with total B7H3 protein (**Figure S2B**).

As AR signaling is a major determinant of ARPC phenotypic subclasses, we determined the relationship of B7H3 RNA and protein to AR target gene output. However, consistent with the analyses of human mPC tumors. there was no correlation of B7H3 protein levels with AR signature scores (**Figure S2C**).

### B7H3-PBD-ADC is cytotoxic for defined subclasses of prostate cancer

We next sought to determine the efficacy of a B7H3 targeted ADC directing the genotoxic PBD (B7H3-PBD-ADC) to mPC cells. We compared the targeted delivery of PBD via B7H3-PBD-ADC relative to the non-targeted control R347-PBD-ADC across a panel of mPC organoids where phenotype, genotype, and B7H3 levels were established (**Figure 2, A and B, Figure S3A, and data file S1**). All non-ARPC models were highly sensitive to the B7H3-PBD-ADC with normalized AUC (nAUC) ranging from 0.2-0.5 and IC50 from 0.03-2.08 ng/ml (**Figure 2C and Table 1**). In contrast, the ARPC models displayed a broad range of responses, with nAUC ranging from 0.3 to 1 and displaying less steep dose response slopes in responders compared to the non-ARPC models (**Figure 2B and Figure 2C**). The relative dose required for cytotoxicity in comparing targeted B7H3-PBD-ADCs and control R347-PBD-ADCs was a minimum of 1000-fold for the most sensitive models, while the majority of models were unaffected by even the highest concentration of R347-PBD (4 mg/ml) (**Figure 2B and Figure S3, B and C**).

**FIGURE 2.**
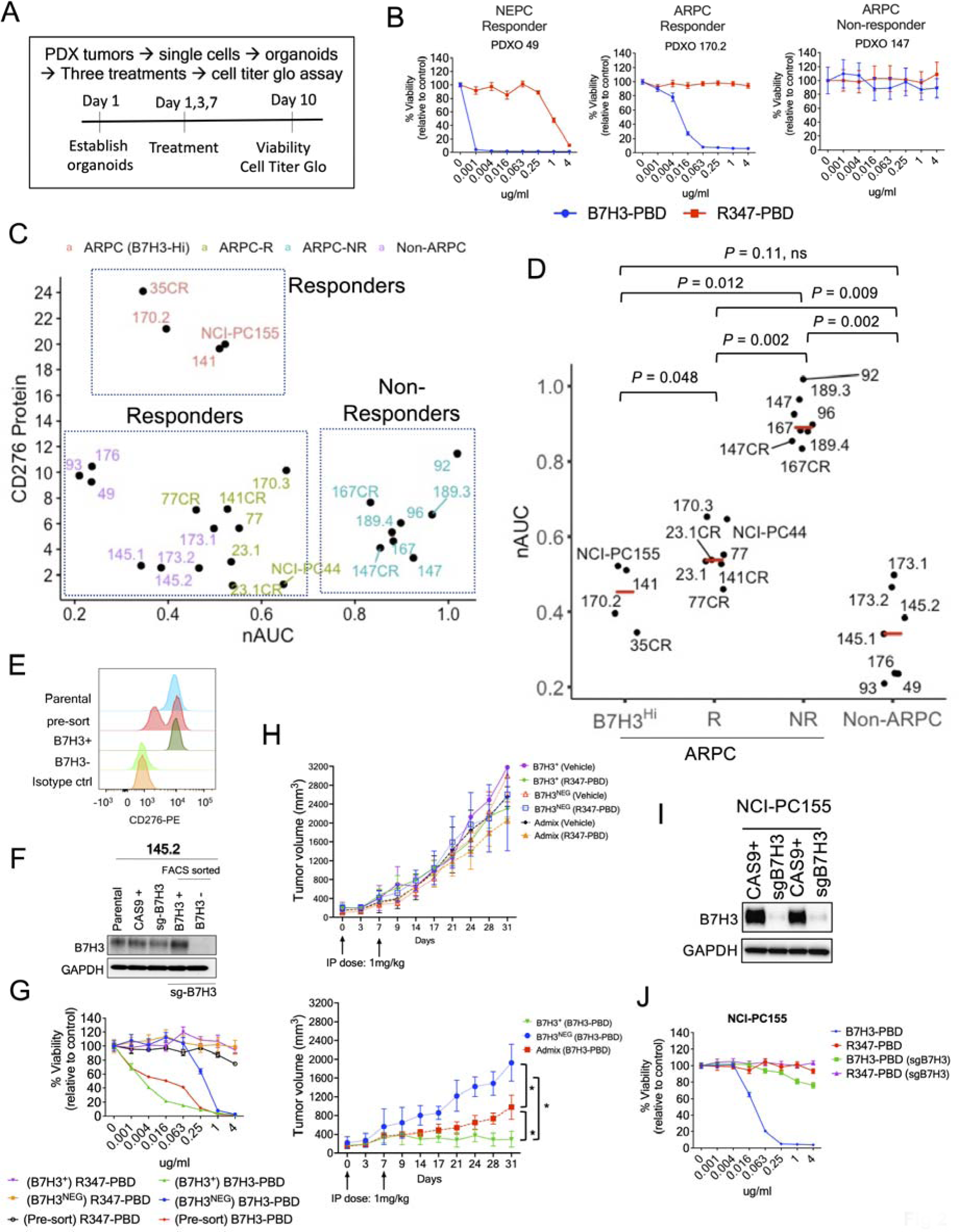
B7H3-PBD-ADC activity requires, but is not correlated with B7H3 protein levels. **(A)** Schematic of the ex-vivo drug assay. **(B)** Representative drug response curves for B7H3-PBD-ADC and R347-PBD-ADC (control ADC) in PDXOs of SCNPC and ARPC phenotypes. % Viability was plotted relative to the control. **(C)** Comparison of B7H3-PBD-ADC response and B7H3 protein expression across the models; n=26. **(D)** Normalized AUC (nAUC) values for B7H3-PBD-ADC in ARPC (n=19) and non-ARPC models (n=7). ARPC models are categorized into three groups: high B7H3 expressors (B7H3^HI^) n=4, responder (R) n=7, and non-Responder (NR) n = 8. Red line indicates median nAUC for the groups. **(E)** FACS sorting strategy for selecting B7H3-KO 145.2 cells. **(F)** Western blot for FACS sorted 145.2 B7H3^+^ and B7H3^-^ (B7H3-KO) cells grown as organoids. **(G)** Dose response curves for 145.2 pre-sorted, and sorted B7H3^NEG^ and B7H3^+^ organoids treated with ADC for 10 days. **(H)** 145.2 B7H3^+^, B7H3^NEG^, and admix (mix of B7H3^+^ and B7H3^NEG^ cells in approximately equal proportion) organoid derived xenografts (ODXs) treated with ADCs or vehicle, once weekly for 2 weeks, as indicated by arrows; n = 8/group, except B7H3^NEG^ (2 mice with necrotic tumors at Day 14 excluded from B7H3-PBD group), B7H3^NEG^ and admix (Vehicle group; n = 2 each), B7H3^+^ (Vehicle group; n = 4), Admix (B7H3-PBD and R347-PBD; n = 5 each). Average tumor volume is plotted from the day of first treatment indicated as Day 0. Top panel comparing average tumor volumes for R347-PBD and vehicle treated mice. Bottom panel comparing average tumor volumes for B7H3-PBD treated B7H3^+^, B7H3^NEG^, and admix xenografts. **(I)** Western blot for B7H3 knockdown in NCI-PC155 organoids. **(J)** Dose response curves for NCI-PC155 organoids after B7H3 knockdown (sgB7H3 group). Error bars indicate the SEM, \**P* < 0.05.

**Table 1.**
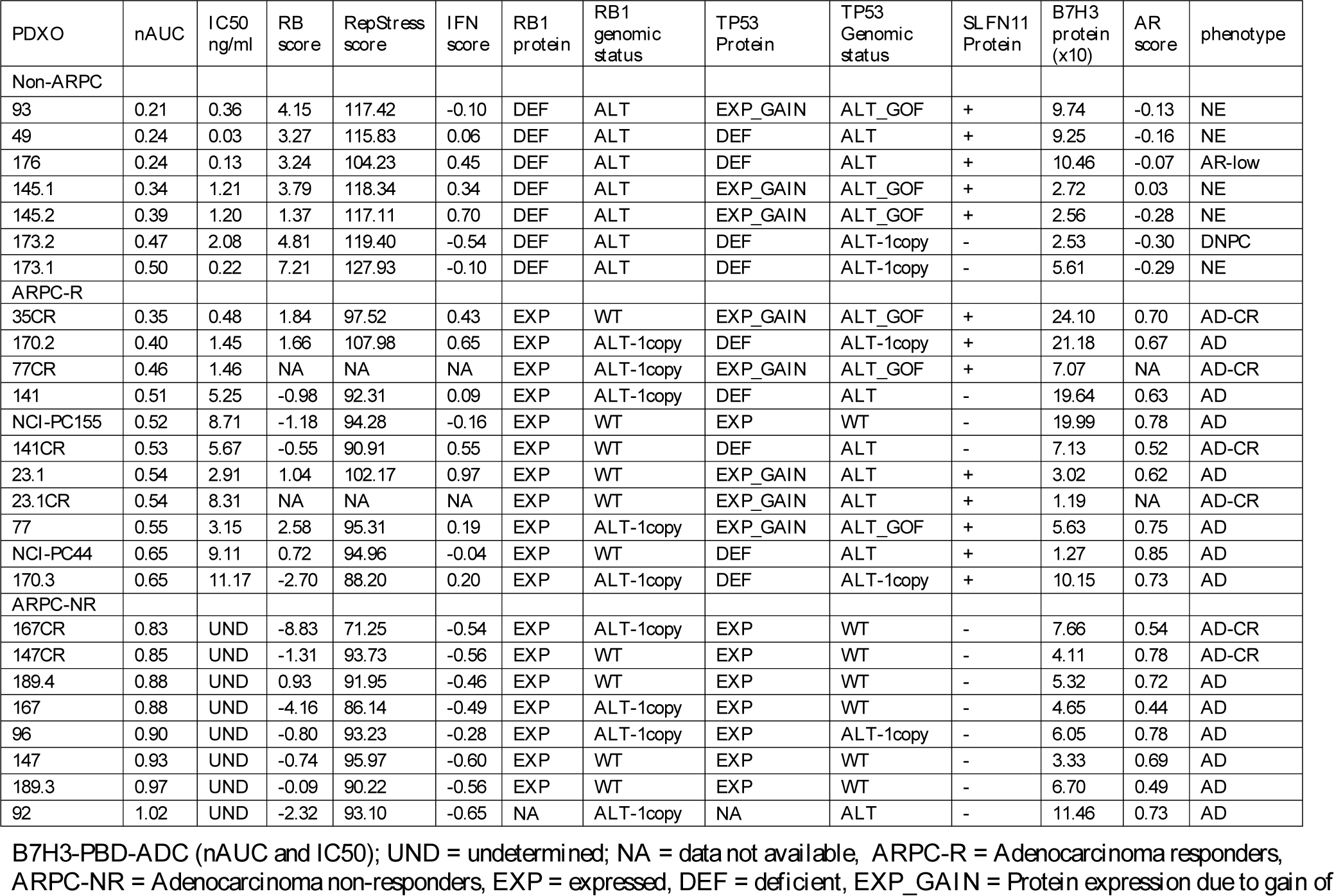
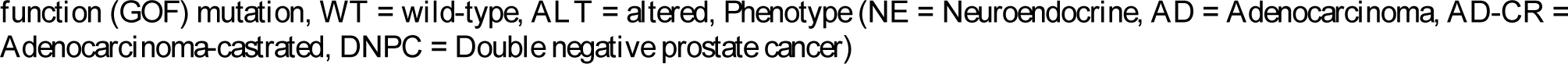
B7H3-PBD response data, and molecular features of 26 mCRPC models.

| PDXO | nAUC | IC50<br>ng/ml | RB<br>score | RepStress<br>score | IFN<br>score | RB1<br>protein | RB1<br>genomic<br>status | TP53<br>Protein | TP53<br>Genomic<br>status | SLFN11<br>Protein | B7H3<br>protein<br>(x10) | AR<br>score | phenotype |
| --- | --- | --- | --- | --- | --- | --- | --- | --- | --- | --- | --- | --- | --- |
| **Non-ARPC** |  |  |  |  |  |  |  |  |  |  |  |  |  |
| 93 | 0.21 | 0.36 | 4.15 | 117.42 | -0.10 | DEF | ALT | EXP_GAIN | ALT_GOF | + | 9.74 | -0.13 | NE |
| 49 | 0.24 | 0.03 | 3.27 | 115.83 | 0.06 | DEF | ALT | DEF | ALT | + | 9.25 | -0.16 | NE |
| 176 | 0.24 | 0.13 | 3.24 | 104.23 | 0.45 | DEF | ALT | DEF | ALT | + | 10.46 | -0.07 | AR-low |
| 145.1 | 0.34 | 1.21 | 3.79 | 118.34 | 0.34 | DEF | ALT | EXP_GAIN | ALT_GOF | + | 2.72 | 0.03 | NE |
| 145.2 | 0.39 | 1.20 | 1.37 | 117.11 | 0.70 | DEF | ALT | EXP_GAIN | ALT_GOF | + | 2.56 | -0.28 | NE |
| 173.2 | 0.47 | 2.08 | 4.81 | 119.40 | -0.54 | DEF | ALT | DEF | ALT-1copy | - | 2.53 | -0.30 | DNPC |
| 173.1 | 0.50 | 0.22 | 7.21 | 127.93 | -0.10 | DEF | ALT | DEF | ALT-1copy | - | 5.61 | -0.29 | NE |
| **ARPC-R** |  |  |  |  |  |  |  |  |  |  |  |  |  |
| 35CR | 0.35 | 0.48 | 1.84 | 97.52 | 0.43 | EXP | WT | EXP_GAIN | ALT_GOF | + | 24.10 | 0.70 | AD-CR |
| 170.2 | 0.40 | 1.45 | 1.66 | 107.98 | 0.65 | EXP | ALT-1copy | DEF | ALT-1copy | + | 21.18 | 0.67 | AD |
| 77CR | 0.46 | 1.46 | NA | NA | NA | EXP | ALT-1copy | EXP_GAIN | ALT_GOF | + | 7.07 | NA | AD-CR |
| 141 | 0.51 | 5.25 | -0.98 | 92.31 | 0.09 | EXP | ALT-1copy | DEF | ALT | - | 19.64 | 0.63 | AD |
| NCI-PC155 | 0.52 | 8.71 | -1.18 | 94.28 | -0.16 | EXP | WT | EXP | WT | - | 19.99 | 0.78 | AD |
| 141CR | 0.53 | 5.67 | -0.55 | 90.91 | 0.55 | EXP | WT | DEF | ALT | - | 7.13 | 0.52 | AD-CR |
| 23.1 | 0.54 | 2.91 | 1.04 | 102.17 | 0.97 | EXP | WT | EXP_GAIN | ALT | + | 3.02 | 0.62 | AD |
| 23.1CR | 0.54 | 8.31 | NA | NA | NA | EXP | WT | EXP_GAIN | ALT | + | 1.19 | NA | AD-CR |
| 77 | 0.55 | 3.15 | 2.58 | 95.31 | 0.19 | EXP | ALT-1copy | EXP_GAIN | ALT_GOF | + | 5.63 | 0.75 | AD |
| NCI-PC44 | 0.65 | 9.11 | 0.72 | 94.96 | -0.04 | EXP | WT | DEF | ALT | + | 1.27 | 0.85 | AD |
| 170.3 | 0.65 | 11.17 | -2.70 | 88.20 | 0.20 | EXP | ALT-1copy | DEF | ALT-1copy | + | 10.15 | 0.73 | AD |
| **ARPC-NR** |  |  |  |  |  |  |  |  |  |  |  |  |  |
| 167CR | 0.83 | UND | -8.83 | 71.25 | -0.54 | EXP | ALT-1copy | EXP | WT | - | 7.66 | 0.54 | AD-CR |
| 147CR | 0.85 | UND | -1.31 | 93.73 | -0.56 | EXP | WT | EXP | WT | - | 4.11 | 0.78 | AD-CR |
| 189.4 | 0.88 | UND | 0.93 | 91.95 | -0.46 | EXP | WT | EXP | WT | - | 5.32 | 0.72 | AD |
| 167 | 0.88 | UND | -4.16 | 86.14 | -0.49 | EXP | ALT-1copy | EXP | WT | - | 4.65 | 0.44 | AD |
| 96 | 0.90 | UND | -0.80 | 93.23 | -0.28 | EXP | ALT-1copy | EXP | ALT-1copy | - | 6.05 | 0.78 | AD |
| 147 | 0.93 | UND | -0.74 | 95.97 | -0.60 | EXP | WT | EXP | WT | - | 3.33 | 0.69 | AD |
| 189.3 | 0.97 | UND | -0.09 | 90.22 | -0.56 | EXP | WT | EXP | WT | - | 6.70 | 0.49 | AD |
| 92 | 1.02 | UND | -2.32 | 93.10 | -0.65 | NA | ALT-1copy | NA | ALT | - | 11.46 | 0.73 | AD |

B7H3-PBD-ADC (nAUC and IC50); **UND** = undetermined; **NA** = data not available, **ARPC-R** = Adenocarcinoma responders, **ARPC-NR** = Adenocarcinoma non-responders, **EXP** = expressed, **DEF** = deficient, **EXP\_GAIN** = Protein expression due to gain of
function (GOF) mutation, **WT** = wild-type, **ALT** = altered, Phenotype (**NE** = Neuroendocrine, **AD** = Adenocarcinoma, **AD-CR** = Adenocarcinoma-castrated, **DNPC** = Double negative prostate cancer)

Interestingly, there were two categories of ARPC responders to B7H3-PBD-ADCs, which segregated with B7H3 protein expression levels. Organoids with the highest B7H3 protein expression (LuCaPs-35CR, 170.2 and 141, and NCI-PC155) were some of the most sensitive among the ARPC models (**Figure 2, C and D**). There were no non-responders among the highest B7H3 expressing ARPC models. However, many responder and non-responder models had similar B7H3 expression suggesting that ADC sensitivity is influenced by additional tumor biological properties. The ARPC cluster with low to medium B7H3 expression (B7H3^LOW^) contained responder (labeled as “R”) and non-responder (labeled as “NR”) models. Notably, the median protein level in the ARPC (B7H3^LOW^) responders was not significantly different from non-responder ARPC models (5.61 vs 5.63, respectively, *P=*0.58) or from responder non-ARPC models (median B7H3 levels = 5.68) (**Figure 2C**).

To establish specificity, we assessed whether B7H3 protein expression is necessary for ADC activity. CRISPR/Cas9 was used to generate B7H3 null organoid cells in LuCaP 145.2 and NCI-PC155 models. Because mPC organoids cannot be cloned, following B7H3 guide transduction into Cas9 expressing organoids, cells were sorted after several generations of growth based upon B7H3 expression (**Figure 2, E and F**). The loss of B7H3 in B7H3^NEG^ LuCaP 145.2 did not result in any discernable difference in growth compared to B7H3^+^ organoids (**Figure S3D**), indicating that B7H3 expression does not affect autonomous growth rate. Consistent with this, dropout screens utilizing 2 separate B7H3-directed guides in LuCaP 145.2 and LuCaP 173.1 organoids demonstrated no growth selectivity across the entire population (**Figure S3E**). Importantly, loss of B7H3 in 145.2 organoids abrogated response to B7H3-PBD-ADC at concentrations less than 0.25 ug/ml in vitro, demonstrating a >400-fold increased IC50 compared to B7H3-WT organoids (**Figure 2G**). However, we observed cytotoxic effects of the B7H3-PBD-ADC at higher concentrations of 1ug/ml and 4ug/ml, probably resulting from B7H3^+^ contaminants in the B7H3^-^ sorted population contributing bystander effects from free PBD (11). Interestingly, organoids derived from pre-sorted cells containing a mix of B7H3^+^ and B7H3^NEG^ cells, representative of intra-tumor heterogeneity, were almost equally sensitive relative as B7H3^+^ organoids, perhaps as a result of combined cell death via B7H3+ specific ADC activity and bystander killing effect (**Figure 2G, red line)**. To validate B7H3-PBD-ADC specificity in vivo, we used organoid derived xenograft models (ODXs) from sorted B7H3^+^ and B7H3^NEG^ LuCaP 145.2 organoids. To investigate the extent of bystander effect in extreme cases of heterogeneity, we also compared ADC activity in xenografts derived after mixing approximately equal proportions of B7H3^+^ and B7H3^NEG^ cells (labeled as “admix tumors”). Consistent with in vitro analysis, vehicle treated B7H3^+^ and B7H3^NEG^ ODXs displayed similar growth rates in vivo (**Figure 2H, top panel**). B7H3^NEG^ ODXs displayed no discernable response to the B7H3-PBD-ADC (bottom panel, blue line), whereas significant tumor regression was observed in B7H3^+^ tumors (bottom panel) compared to vehicle treated or negative control R347-PBD-ADC treated mice (top panel) (**Figure 2H**). Interestingly, admix tumors showed partial regression, although the tumor growth was significantly slower than B7H3^NEG^ ODXs after the B7H3-PBD-ADC treatment (**Figure 2H, bottom panel**). Immunohistochemical (IHC) analysis of tumors collected at the end of the study showed that B7H3+ cells were eradicated from both B7H3^+^ and admix tumors (**Figure S3, F and G**), demonstrating specificity and suggesting that a proportion of target negative tumor cells escape by-stander mediated killing. Finally, loss of B7H3 protein in NCI-PC155 patient derived adenocarcinoma organoids substantially reversed B7H3-PBD-ADC sensitivity (**Figure 2, I and J**). These results demonstrate that the B7H3-PBD-ADC is specific for B7H3 expressing mPC organoids over a broad range of tested concentrations.

### B7H3-PBD-ADC response is associated with *RB1* deficiency and replication stress

An inspection of the model genotypes relative to B7H3-PBD-ADC efficacy to identify predictive response characteristics revealed that combined alterations of *RB1* and *TP53* were strongly correlated with responsiveness to B7H3-PBD-ADC treatment. Indeed, *RB1* homozygous deletion and *TP53* alterations, either deletion or mutation, occur in all the non-ARPC models (**Figure 3, A and B, and Table 1**). Further, *RB1* function was analyzed across the models using an RB signature score that captures transcriptional networks related to RB1 functional inactivation. The RB signature score was linearly related to B7H3-PBD-ADC sensitivity as measured by AUC in organoid models, suggesting not only a categorical relationship but also a quantitatively determined sensitivity (**Figure 3C**).

**FIGURE 3.**
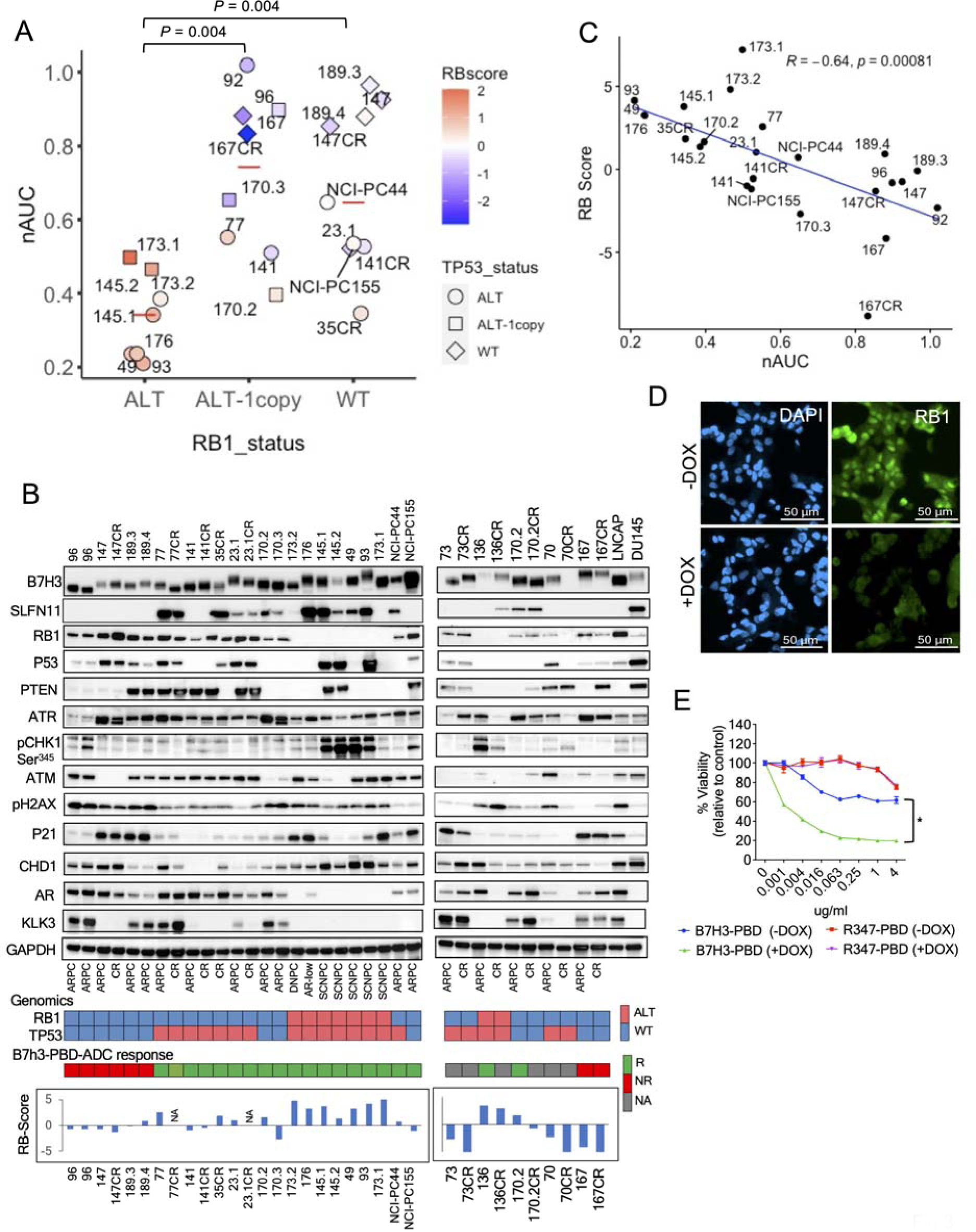
RB1 loss predicts B7H3-PBD-ADC response. (A) B7H3-PBD-ADC response categorized by *RB1* genomic status. Red line indicates the median nAUC for each group. *TP53* genotypes are shown as different shapes. Color indicates RB1 signature score on z-transformed scale. (B) Immunoblot analysis of organoid models and prostate cancer cell lines for the indicated markers. Heatmap (bottom) showing *RB1* and *TP53* genomic status, and B7H3-PBD-ADC response. For *RB1*, red color indicates biallelic copy loss and blue indicates wild-type or 1 copy loss. *TP53* status in red refers to alterations by bi-allelic inactivation or gain of function mutation and in blue indicates wild-type or monoallelic loss. For B7H3-PBD-ADC response, “R” = responsive, “NR” = non-response, “NA” = Data not available. Bar plots for RB1 score is shown for the organoid models. (C) Correlation of B7H3-PBD-ADC response (nAUC) and RB1 signature score. Pearson’s correlation coefficient r = −0.64, *P* = 0.00081. (D) IF images confirming DOX-inducible knockdown of RB1 in LuCaP167 organoid model. (E) B7H3-PBD-ADC dose response curves in LuCaP167 (RB1^+^) organoid model expressing DOX inducible *RB1* shRNA. Error bars indicate the SEM. \**P* < 0.05.

We experimentally validated *RB1* levels as a determinant for B7H3-PBD-ADC responses by depletion via doxycycline-induced shRNA in the non-responsive, *TP53*^WT^*RB1*^WT^ LuCaP 167 ARPC organoid model (**Figure 3D and Figure S4A**). Consistent with the results across the various models, *RB1* depletion resulted in increased B7H3-PBD-ADC cytotoxicity in LuCaP 167 organoids, demonstrating *RB1* levels as an independent factor in determining B7H3-PBD-ADC sensitivity (**Figure 3E**).

Since models with *RB1* loss showed exceptional response to the B7H3-PBD-ADC despite having low levels of B7H3 protein, we also tested the influence of this underlying genomic vulnerability on the in vitro activity of free PBD dimer. Indeed, increased sensitivity to free PBD dimer was observed in *RB1* loss models compared to *RB1*^WT^ models, suggesting that the genotoxic effect of PBD induced inter-strand DNA crosslinks is potentiated by loss of functional *RBI* (**Figure S4B**). Thus, the response to the B7H3-PBD-ADC is substantially governed by the interplay between tumor characteristics and the mechanism of action for the attached payload and not necessarily by the density of the targeted antigen.

*RB1* loss is associated with replication stress, and not unexpectedly, replication-stress induced pathways also correlated with the strength of B7H3-PBD-ADC responses. We used a replication stress signature score (RepStress score), modified for prostate cancer based on DNA damage and cell cycle pathways involved in replication stress, and analyzed association with B7H3-PBD sensitivity (**Figure 4A**). Across all models, including ARPC and non-ARPC groups, a high RepStress score was significantly correlated with higher sensitivity to B7H3-PBD-ADC (**Figure S4C**), a finding which is strongly determined by loss of RB functionality (**Figure S4D**). Moreover, B7H3-PBD-ADC showed greater efficacy in vitro, in comparison with other known Repstress sensitive drugs including Topotecan, Cisplatin, and Mitomycin C (**Figure S4, E and F)**. For the ARPC models only, the majority of the ARPC responders (“ARPC-R”) had similar replication stress scores to non-responders (“ARPC-NR”) with the exception of two responder RB1^WT^ models (LuCaPs 23.1 and 170.2) that demonstrate RepStress scores above the average (**Figure 4B)**. This implies that PBD sensitivity of mPC with functional RB1, usually ARPC, is determined by factors in addition to replication stress.

**FIGURE 4.**
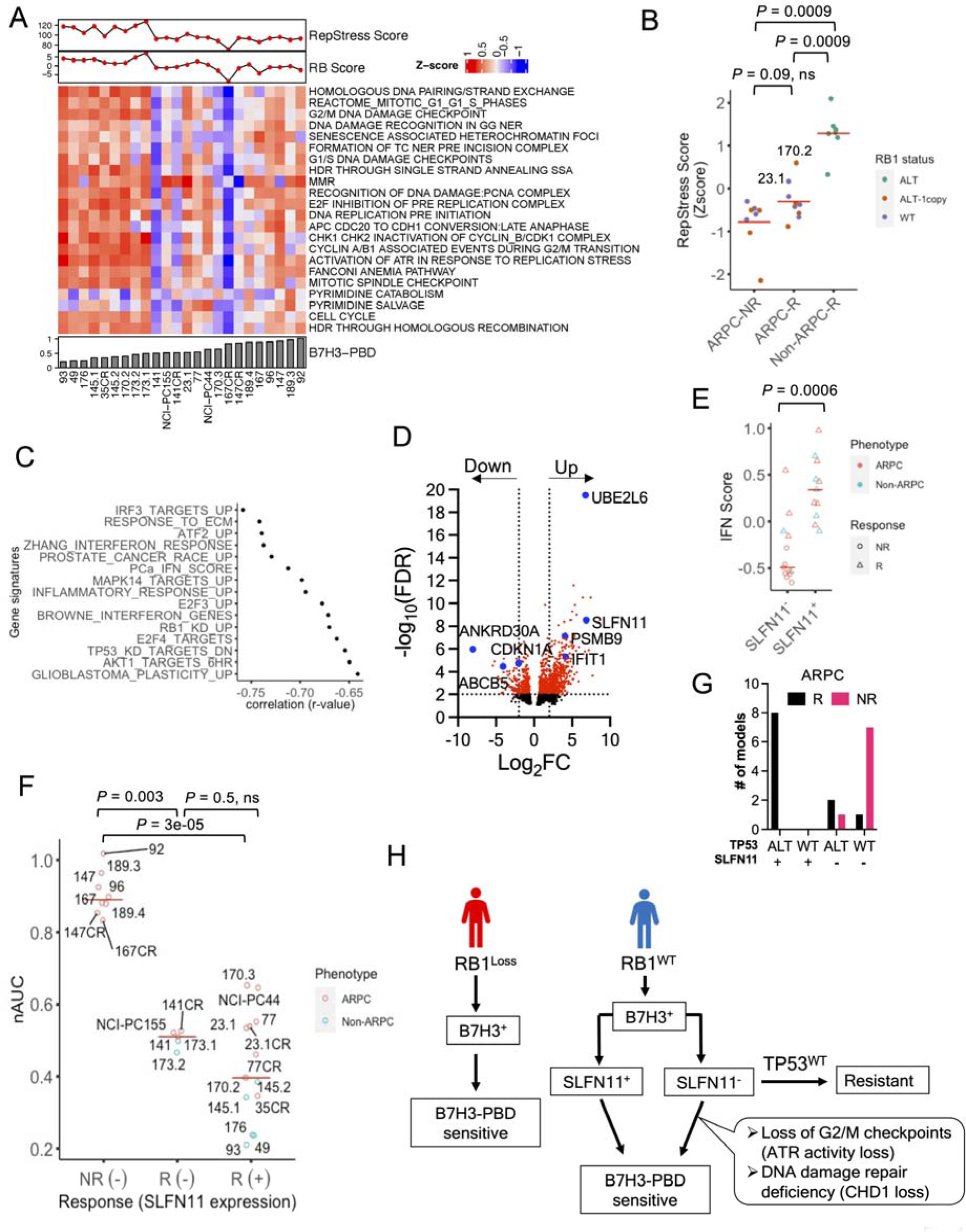
Contributing biomarker subclasses of B7H3-PBD-ADC sensitivity. **(A)** Heatmap of the pathways contributing to the replication stress (RepStress) signature score. Organoid models are ranked from left to right based on increasing B7H3-PBD nAUC (bottom panel). Top panel showing RepStress and RB signature scores. **(B)** Comparison of z-transformed RepStress score in AD non-responders (ARPC-NR), ARPC responders (ARPC-R), and non-ARPC responders (NonARPC-R). Color indicates *RB1* genotype. **(C)** Univariate correlation analyses between nAUC and MsigDB gene signatures, including refined IFN signature score for prostate cancer labeled as “PCa_IFN_Score”. Spearman correlation coefficient is shown for top significant gene sets. FDR ≤ 0.05. **(D)** Volcano plot for differentially expressed genes between B7H3-PBD-ADC responsive ARPC models vs non-responsive ARPC models. Dotted lines are shown for log_2_ fold change of −2 and 2 at FDR ≤ 0.01. **(E)** Comparison of IFN score with SLFN11 expression. **(F)** B7H3-PBD-ADC response categorized by SLFN11 expression. Red line indicates median nAUC for each group. NR = non-responder, R = responder, +/-indicates SLFN11 expression. **(G)** Distribution of ARPC models based on *TP53* genomic status and SLFN11 expression**. (H)** Schematic of proposed biomarker based therapeutic decisions for B7H3-PBD-ADC treatment of mPC patients.

### SLFN11 expression and TP53 status are predictors of B7H3-PBD-ADC response in RB1 functional prostate cancer

To address broadly predictive biomarkers in ARPC with functional *RB1*, the most common clinical phenotype of mPC, we analyzed differentially expressed signaling pathways in responder and non-responder ARPC models (ARPC-R vs ARPC-NR). Correlation analysis with single sample gene set enrichment scores identified interferon response gene signatures as the topmost correlated signaling pathways for B7H3-PBD-ADC sensitivity (nAUC) for ARPC models (**Figure 4C**). To further delineate the role of specific interferon stimulated genes (ISGs) that may be contributing to B7H3-PBD-ADC response, we performed differential expression analyses comparing ARPC responders (n = 9) and non-responders (n=8), which revealed several ISGs among the top 20 differentially upregulated genes in responders including *UBE2L6*, *PSMB9*, and *SLFN11* while *CDKN1A* and *ABCB5* were notably downregulated (**Figure 4D and data file S2**). Upregulation of SLFN11 in ARPC responder models was of particular interest as it is a known sensitizer for toxicity mediated by specific DNA damaging agents (22, 23). *SLFN11* is a non-classical interferon (IFN)-response gene, and indirect effects of IFN signaling likely contribute to contextual SLFN11 expression (22). As expected, in the phenotypically heterogeneous models analyzed here, SLFN11^NEG^ models demonstrated a significantly lower median IFN signature score than SLFN11^+^ models (**Figure 4E**). We categorized B7H3-PBD-ADC response based on SLFN11 positivity or negativity. Consistent with differential expression analysis, SLFN11 expression predicted response in 8 of 8 ARPC models, demonstrating SLFN11 as a robust positive biomarker in the models analyzed here (**Figure 4F)**. Of significance, we observed that enrichment for wild-type TP53 alleles was a common molecular characteristic amongst 7/8 SLFN11^NEG^ non-responders, suggesting that SLFN11 expression may be linked to TP53 mutation status (**Figure 4G and Table 1**). It should be noted that a small number of SLFN11^NEG^ models were also responsive to the ADC (3 responders out of 11 SLFN11^NEG^ ARPC models) suggesting that lack of SLFN11 expression is not always predictive and that other non-overlapping mechanisms/biomarker classes also lead to sensitivity **(Figure 4F,R)** in ARPC, as described below. Similarly, the majority (6/9) of RB1 deficient models were SLFN11^+^ (**Figure 3B**), but RB1 deficient models were responsive to B7H3-PBD-ADC independent of SLFN11 expression, demonstrating that replication stress predicts sensitivity even in the absence of SLFN11 (**Figure 4F, R**(**-**) **group; 173.1 and 173.2)**.

In B7H3 expressors*, RB1* loss/ replication stress and/or SLFN11 expression predicted the responses of most tumors to B7H3-PBD-ADC treatment, but there were outliers where these features did not discriminate outcomes. For example, LuCaPs 141 and 141CR, and NCI-PC-155 were categorized as *RB1*^WT^ and SLFN11^NEG^ responders (**Figure 3B and Figure 4G**). We analyzed known genomic vulnerabilities associated with inability to repair inter-strand crosslinks and expression of previously known transporters of PBD. None of the previously identified transporters of PBD (*ABCG2, ABCB1, ABCC2, and SLC46A3*) correlated with responsiveness in the extensive mPC cohort tested here (**Figure S4G**). Because ATR loss of function is a sensitizing factor for PBD responsiveness (24), we performed ATR activation assay in response to DNA damage in ARPC organoids of B7H3-PBD-ADC responders: LuCaP 77 (*RB1*^WT^, SLFN11^+^), LuCap 141CR and NCI-PC155 (*RB1*^WT^, SLFN11^NEG^) and the non-responder LuCaP 167 (*RB1*^WT^, SLFN11^NEG^). Functional ATR was evident in response to topotecan and B7H3-PBD-ADC in the responder LuCaP 77 and LuCaP141CR models. However, ATR was reduced to near undetectable levels in responder NCI-PC155 and non-responder LuCaP 167 (**Figure S5, A-D**). Consistent with non-responsiveness, LuCaP 167 showed no evidence of PBD-mediated DNA damage. By contrast, NCI-PC155 had clearly detectable topotecan and B7H3-PBD-ADC induced DNA damage (gH2AX) compared to LuCaP 167 (**Figure S5, A-B**). These data for NCI-PC155 are consistent with a mechanism whereby very weak ATR signaling enhances DNA damage-induced death.

As the SLFN11^NEG^ LuCaP 141CR model had an intact ATR pathway, we considered other DNA repair biomarkers. Importantly, we observed loss of CHD1 protein in LuCaPs 141 and 141CR (**Figure 3B**). CHD1 deficient cells are generally hypersensitive to DNA cross-linking agents because of defects in homologous recombination mediated repair, and in clinical mPC, *CHD1* mutations are statistically associated with a predicted homologous recombination deficiency (HRD) (25, 26). These data are consistent with a responsive phenotype due to a DNA repair deficiency for LuCaPs 141 and 141CR despite an *RB1^WT^*genotype and lack of SLFN11 expression (**Figure 4H**).

Indeed, our analysis of ATR activity and CHD1 loss is limited by the number of available models, which are derived from CRPC patient populations in which *ATR* and *CHD1* mutations occur at a frequency of < 5%. Although the proposed mechanisms for responsiveness in these *RB1^WT^*SLFN11^NEG^ models require validation, the observation of their existence is notable and directs future studies to investigate relatively infrequent DNA repair deficiency-mediated responsive mechanisms.

### In vivo models of mPC validate organoid response classes to B7H3-PBD-ADC therapy

Based on the in vitro B7H3-PBD-ADC response data and analysis of potential biomarkers, we evaluated the efficacy of B7H3-PBD-ADC treatment in preclinical trials of PDX models representative of different phenotype and genotype mPC categories defined by organoid studies: *RB1*^NULL^ (SLFN11^+^ or SLFN11^-^) and *RB1*^WT^ (SLFN11^+^ or SLFN11^-^) tumors (**Figure 5**). We randomized mice implanted with PDX lines to treatment with two intraperitoneal doses of 1mg/kg B7H3-PBD-ADC or control R347-PBD-ADC, given weekly for two weeks. The SCNPC LuCaP 145.2 (*TP53*^ALT^*RB1*^-/-^SLFN11^+^) line showed a complete and durable response to B7H3-PBD-ADC treatment (**Figure 5A**). Control R347-PBD-ADC and vehicle-treated mice had similar tumor growth indicating no apparent non-specific effects of PBD at a 1 mg/kg dose. Remarkably, no tumors were detected in 8 out of 9 B7H3-PBD-ADC treated mice 3 months after therapy. Furthermore, two of the large established tumors (>1000 mm3) were completely regressed with just two doses of B7H3-PBD-ADC (**Figure 5A, right panel**).

**FIGURE 5.**
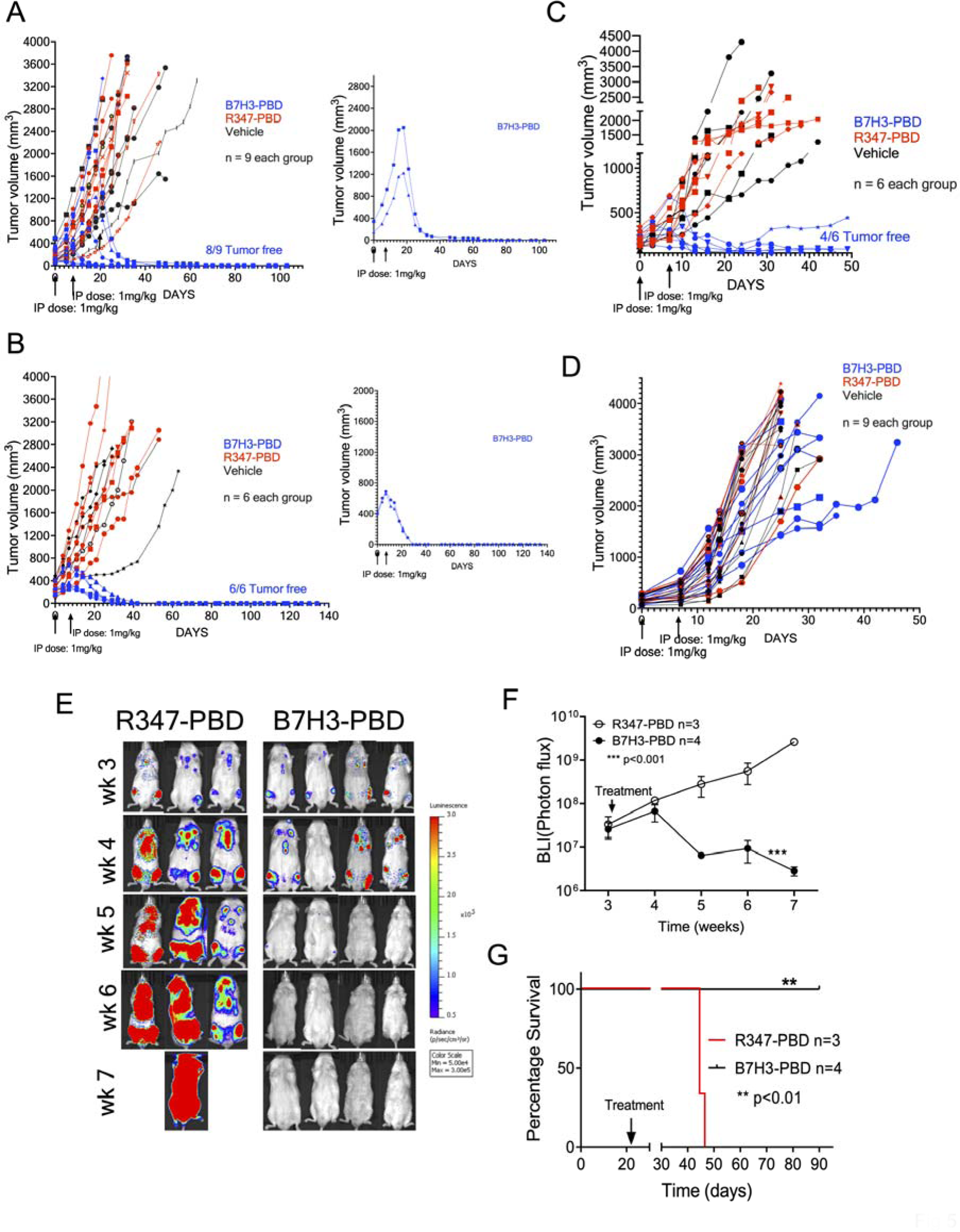
Prostate cancer organoid-derived biomarkers predict in vivo tumor responses in preclinical trials of the B7H3-PBD-ADC. (**A-D)** Tumor response to B7H3-PBD-ADC (1mg/kg), R347-PBD-ADC (1mg/kg) or vehicle in the four selected LuCaP models based on identified biomarkers **(A)** LuCaP 145.2 (SCNPC phenotype; RB1^loss^, SLFN11^+^, IFN score^HIGH^), n=9 / group. **(B)** LuCaP 136 (ARPC phenotype; RB1^loss^, SLFN11^-^, IFN score^low^), n = 6/ group. **(C)** LuCaP 77 (ARPC, RB1^+^, SLFN11^+^, IFN score^medium^) n = 6 /group. **(D)** LuCaP 167 (ARPC, RB1^+^, SLFN11^-^, IFN score^low^) n = 9 /group. Right panels for (A) and (B) displaying antitumor activity of B7H3-PBD-ADC in mice with large established tumors (145.2; >1000 mm^3^ and 136; >650 mm^3^). Tumor volume measurements (mm^3^) are shown from the time of first treatment. Arrows indicate once weekly dose for 2 weeks. **(E)** Bioluminescence imaging of LuCaP 136 metastases following treatment with B7H3-PBD-ADC (n = 4) or R347-PBD-ADC (n=3). Mice were treated once weekly for two weeks. **(F)** Average BLI for treated mice from the time of first treatment. Means ± standard deviation, Two-way ANOVA test; *P* < 0.001. (G) Kaplan-Meier survival analysis for the B7H3-PBD-ADC and R347-PBD-ADC treated LuCaP 136 metastases. Log-rank test (*P* < 0.01).

Similarly, the LuCaP 136 ARPC PDX (*TP53*^-/-^*RB1*^-/-^SLFN11^-^) showed complete durable response in all 6 mice treated with B7H3-PBD-ADC compared to control R347-PBD-ADC treated or vehicle treated mice (**Figure 5B**). LuCaP 136 was not evaluable for in vitro responses due to poor growth characteristics beyond 1 week, but the *RB1* loss phenotype predicted in vivo responsiveness. Again, B7H3-PBD-ADC treatment resulted in a remarkable decrease in large tumor burden (∼800 mm^3^) in 2 mice (**Figure 5B, right panel**). 100% of the mice were tumor free for more than 4 months after treatment. Further, B7H3-PBD-ADC treatment of the ARPC LuCaP 77 xenograft (*RB1*^WT^SLFN11^+^) also showed a durable response relative to R347-PBD-ADC treated mice (**Figure 5C**). In contrast, the ARPC LuCaP 167 xenograft (*RB1*^WT^SLFN11^-^) which did not respond to B7H3-PBD-ADC in vitro, also showed no significant tumor regression with B7H3-PBD-ADC treatment compared to R347-PBD-ADC treated or vehicle treated mice (**Figure 5D**). Thus, the treatment responses assessed via organoid assays accurately predicted the matching in vivo responses to B7H3-PBD-ADC.

Because the majority of clinical mPCs progress following ADT and ARSI treatment with the retention of AR signaling, and patients often harbor bone metastases, we further tested B7H3-PBD-ADC activity against a late-stage ARPC bone metastasis model. We developed a model system whereby intracardiac injection of luciferase-tagged AR^+^ LuCaP 136 (*TP53*^-/-^*RB1*^-/-^ SLFN11^-^) tumor cells colonized bone with 100% efficiency. The majority of metastases were located in the vertebrae. Other sites included calvaria, sternum, and long bones, as well as <10% of tumor burden in soft tissue (adrenals and liver). Using bioluminescence imaging (BLI) to monitor the growth of tumor metastasis, mice were grouped into two arms of equal average metastasis burden. B7H3-PBD-ADC treatment once weekly, for two weeks, substantially reduced tumor burden resulting in no detectable metastasis compared to control R347-PBD-ADC treated mice (**Figure 5, E and F**). Overall, B7H3-PBD-ADC treatment resulted in long-term metastasis-free survival in 100% of the treated mice (**Figure 5G**). These data extend B7H3-PBD-ADC efficacy to tumors residing in clinically significant microenvironments.

We evaluated the safety profile of B7H3-PBD-ADC both in vitro and in vivo. Body weights were unaffected by the ADC in all in vivo preclinical trials (**Figure S6, A-D**). In addition, we performed full necropsy on day 8 and day 30 post-treatment to evaluate acute and delayed in vivo toxicity, respectively, in LuCaP 136 PDX model after administrating the ADC (n= 3 per group). Histopathology analysis demonstrated a reasonable safety profile with no obvious changes in gross pathology. Hematoxylin and eosin-stained sections of liver, heart, lung, brain, adrenal gland, kidney, small intestine, spleen, and prostate collected on day 8 or day 30 post-treatment from all animals were examined. Treatment related microscopic changes were limited to minimal small intestine crypt apoptosis in R347 or B7H3 treated animals on day 8 post-treatment (**Figure S6E**). This change had recovered by day 30 post-treatment. All remaining microscopic changes were similar between untreated and treated animals or consistent with common background findings in mice. Further, human IgG immunoreactivity was not observed in any study tissues examined from vehicle, R347-PBD-ADC or B7H3-PBD-ADC treated animals collected at day 8 or day 30 post-treatment, suggesting no accumulation of ADC in normal mouse tissues. Additionally, we tested the B7H3-PBD-ADC and free PBD payload in organoids derived from normal mouse prostate and normal human liver cells (21). Consistent with the in vivo results, the ADC showed no toxicity in human liver cells **(Figure S6F)** or normal mouse prostate organoids **(Figure S6G)** whereas free PBD affected viability of normal organoids only at significantly higher concentrations (IC50 = ∼2-5nM) than that observed for responder tumor models (IC50 < 0.03 pM) **(Figure S6G and Figure S4B)**.

### Distribution of biomarkers in clinical samples

We related the results presented here (**Figure 4H**) to the distribution of biomarkers in clinical CRPC by analyzing the SU2C mPC data set, considering samples with >30% tumor content (303 samples were included) (**Figure S7**). Within these samples ∼10% demonstrated homozygous *RB1* alterations (27, 28). The influence of SLFN11 upon drug sensitivity has been associated qualitatively with presence of the RNA and protein (19, 23, 24). From the SU2C RNAseq data, we estimated that about 40% of *RB1* intact samples expressed SLFN11 (see methods) (**Figure S7**). Further, of the remaining *RB1*^WT^/*SLFN11*^NEG^ samples (n=153), about 4% of patients (6 of 153) demonstrated CHD1 or ATR loss whereas 91 out of 153 had *TP53*^WT^ and no ATR/CHD1 alterations. In summary, considering any one biomarker as predictive for response, approximately 50% of the SU2C cohort could have been considered further for B7H3-PBD-ADC treatment pending the determination of B7H3 protein expression.

## DISCUSSION

Here we present one of the first examples of biomarker stratification within mPC patient tumors using an organoid/PDX platform exemplifying the extensive genomic and phenotypic heterogeneity of clinical disease. We evaluated the rapidly developing therapeutic modality of antibody-directed cytotoxics, specifically B7H3-targeted PBD, a DNA interstrand cross-linking agent. Novel approaches to treatment of advanced castration resistant mPC are needed as acquired resistance to ARSIs occurs with near universal frequency. We determined that B7H3 is expressed across a diverse spectrum of late-stage treatment-resistant metastatic mPC genotypes and phenotypes as well as across metastatic sites, which extends prior analyses of B7H3 in prostate cancer (7–9, 29).

B7H3 expression is necessary for targeted delivery and ADC response as demonstrated by the loss of responsiveness upon CRISPR/CAS9 mediated B7H3 loss (**Figure 2, E-J**). Importantly, however, B7H3 expression alone is not predictive, as several B7H3^+^ adenocarcinomas were resistant to treatment. Here and with other ADCs, response biomarkers are required to stratify patients to optimize efficacy and minimize ineffective drug exposure. PBD delivered via B7H3-ADC directed exposure exhibited a range of in vitro and in vivo activity against mPC organoid models that reflect underlying biology described by genotypic and phenotypic markers.

Molecular interrogation revealed biomarker-defined classes of responsive models including: 1) *RB1/TP53* loss of function and associated replication stress observed predominantly in SCNPC and in some highly aggressive adenocarcinomas, 2) SLFN11 expression observed in a subset of *RB1* WT/*TP53* altered adenocarcinomas, and 3) specific DNA repair mutations (including *ATR* and *CHD1)* that impact PBD-initiated DNA damage. Thus, multiple vulnerabilities distinct from AR dependent survival and including but expanded beyond homologous recombination deficiencies targeted with PARP inhibitors can be exploited using PBD based therapeutics. The mPC PDX/organoid cohort analyzed here approximately replicates the distribution of identified baseline biomarkers, *RB1* and *TP53* mutations as well as SLFN11 expression, observed in the large SU2C CRPC clinical data set. We anticipate that the classification and associated rationale for biomarker-identified organoids here largely reflects the potential response spectrum in clinical mPC and will inform the design and impact the overall efficacy of prospective trials much more accurately than prior therapeutics analyses with limited numbers of in vitro growth selected mPC cell line models previously available.

A category of biomarkers associated with B7H3-PBD-ADC responses involve a vulnerability to replication stress. A unique conclusion of this study is that mPCs with functional alterations in *RB1/TP53* are highly sensitive to the B7H3-PBD-ADC irrespective of their histological phenotype. Defects in *RB1* and *TP53* pathways are known to promote enhanced replication stress (28), and additional blockage of replication forks by PBD induced ICLs likely contributes to its potent cytotoxicity. It will be useful to determine whether ADCs with PBD payloads have the potential to be efficacious in other cancer types with high replication stress phenotypes.

Although clinical studies have demonstrated that *TP53/RB1* deficient SCNPCs as well as small cell lung cancer generally show high response rate to platinum chemotherapy, many patients respond for relatively short periods of time, and patients almost universally succumb to recurrent disease (32, 33). Here, *TP53/RB1* deficient models of various histological types demonstrated highly efficacious complete responses to B7H3-PBD-ADC. This suggests that B7H3-PBD-ADC may be considered for treatment of *RB1* deficient cancers following platinum resistance.

SLFN11 expression was positively predictive of sensitivity to B7H3-PBD-ADCs. SLFN11 is a DNA/RNA helicase that is actively recruited to sites of DNA damage and appears to irreversibly block stressed replication forks, leading to the hypothesis that SLFN11 is a dominant inhibitor of stressed replication fork repair. SLFN11 is likewise a useful biomarker for response to a variety of other therapeutics that target enhanced replication stress, including topoisomerase and PARP inhibitors as well as platinum chemotherapeutics (19, 23). Of note for the present study of mPCs, we found that a cell-autonomous IFN score was positively associated with expression of SLFN11, a non-canonical IFN response gene (22), and negatively correlated with the presence of a WT *TP53* allele. Future studies to establish the mechanistic basis of this observation may prove of significance for using multiple associated biomarkers to optimize therapeutic predictions and to gain insight into the physiological context of SLFN11 activity.

The direct or indirect repair of PBD-initiated DNA interstrand crosslinks, a major consequence of which is the deleterious blockage of replication forks, is an anticipated class of mechanistic biomarkers. Although further confirmatory studies are required, we present preliminary data that loss of CHD1 appears to sensitize toward B7H3-PBD-ADCs killing, as observed for the SLFN11^NEG^/*RB1*^WT^ adenocarcinoma model, LuCaP 141. By contrast, the LuCaP 96 model, which displays loss of function for *BRCA2,* a classical HRD protein, was non-responsive to B7H3-PBD-ADCs. It is likely significant that LuCaP 96 expressed WT *TP53*, and previous analyses of *BRCA1/2* deficient breast cancer PDXs have suggested approximately 25% are non-responsive to PBD, a phenotype that may be associated with a less compromised HRD resulting from minimal mutations to additional DNA repair proteins, including *TP53* (30, 31). In addition, we characterized a naturally occurring, PBD-responsive, SLFN11^NEG^ *TP53*^WT^ ATR loss of function model, NCI-PC155. This observation is from a natural patient derived model, providing further support for the conclusions drawn from experimentally-selected SLFN11^NEG^ tumor models where inhibition of ATR was synthetically lethal (19, 24).

In summary, B7H3-PBD-ADCs deliver a potent ICL agent with substantial anti-tumor effects toward subtypes of mPC refractory to standard of care treatment regimens. Several acquired resistance mechanisms in mPC appear sensitive to the action of PBD, supporting further evaluation as a therapeutic strategy for cancers that are refractory to currently approved agents. Collectively, our analyses suggest that mPCs vulnerable to PBD action can be identified by a composite set of biomarkers that address multiple histological phenotypes, including adenocarcinomas and SCNPC, and utilize genomic markers (*RB1, TP53, CHD1,* and *ATR)* and/or phenotypic markers (B7H3, SLFN11, and RepStress scores). The approach reported here for identifying biomarkers of vulnerability to a particular cancer therapeutic using an extensive cohort of diverse models is one of the first such studies to be done with metastatic prostate cancer and has the potential to increase the reliability of translation to the clinic as well as to provide insights into mechanisms that underlie the more precise allocation of therapy.

## MATERIALS AND METHODS

### Study design

The objective of this study was to evaluate B7H3 expression and treatment efficacy of the B7H3 ADC armed with a pyrrolobenzodiazepine warhead (SG3315) in a diverse spectrum of PDX/organoid preclinical models of mPC. An additional objective was to identify predictive biomarkers of B7H3-PBD response. In total, 37 mPC organoid models were evaluated for B7H3 expression. At least 26 of 37 organoid models that can be grown in vitro for a minimum of 10 days were tested with the ADCs (**Figure S3A**). Gene expression data for 24 out of 26 organoid models (not available for 23.1CR and 77CR) was utilized to identify correlates of PBD response.

For in vivo studies, the sample size for each experiment is indicated in the figure legend. The animal caretaker who assessed and treated animals, and measured tumors was blinded to the intervention. All other investigators were not blinded for any experiments.

### Properties of anti-B7H3 ADC

B7H3-PBD-ADC is composed of a Human IgG1 anti-B7H3 antibody, site specifically conjugated via a cathepsin-cleavable valine-alanine (val-ala) linker to PBD dimer warhead SG33115 (patent WO 2015/052322). Drug antibody ratio (DAR) was 2. Anti-B7H3 antibody (see supplementary methods for details) is non-mouse cross reactive (**also shown in Figure S6F**) and binds to huB7H3 but not huB7H4 (∼29% homology among the closest family homologs).

### RNA Sequencing Analysis of mPC tumors

RNA isolation and sequencing of 185 UW mPC tumors from 98 patients were performed as described previously (2). Sequencing reads were mapped to the hg38 human genome using STAR.v2.7.3a. All subsequent analyses were performed in R. Gene level abundance was quantitated using GenomicAlignments and transformed to log_2_ FPKM. Groups were compared using two-sided Wilcoxon rank tests with Benjamini-Hochberg multiple-testing correction. To evaluate the expression of B7H3 from tumors within the same patient, boxplots of transcript levels from tumors from the same patient were filtered to include only patients with at least two tumors profiled (149 tumors from 62 patients) and are ordered by per-patient median log_2_ FPKM gene expression.

### Immunohistochemistry

Tissue microarray (TMA) slides consisting of triplicate cores from 181 metastatic prostate sites representing 58 donors was provided by the GU Lab at the University of Washington. Automated IHC was performed on the VENTANA Discovery Ultra (Ventana Medical Systems Inc) autostainer. Details of the method for immunohistochemistry staining are provided in the supplementary methods.

The TMA slides were scanned at a 40X magnification using the Ventana DP 200 instrument (VMSI) and visualized with HALO (Indica Labs). Staining intensity was evaluated by a pathologist (M.R.). The H-score was determined by multiplying the percentage of positive cells at each intensity level (0 denotes no staining, 1 denotes weak staining, 2 denotes moderate staining and 3 is strong staining) by its respective intensity level. The weighted scores were totaled, resulting in a composite score ranging from 0-300. The triplicate scores for each site were averaged to generate an averaged H-score for each site.

### Organoid culture

LuCaP PDX tissue samples were processed into single cells, depleted for mouse cells using mouse cell depletion kit (Miltenyi Biotec, manufacturer’s protocol), and plated as organoids according to our previously described methods (21). Organoids were grown in 5% Matrigel on ultra-low attachment plates/dishes (Corning) for short-term drug treatment and flow cytometry analysis. Previously defined organoid culture media (21) was supplemented with 5ng/ml Neuregulin 1 (Peprotech, 100-03) to potentiate the growth of adenocarcinoma organoid models.

### Pharmacological agents

Two batches of antibody drug conjugates (B7H3-PBD or R347-PBD) were received from AstraZeneca. Organoid responses with an individual batch of the ADC are shown in Figure S3A. No major difference in drug response was observed. The R347 is a control human IgG1 antibody that does not bind human proteins and serves as the isotype control antibody drug conjugate. Free PBD payload was purchased from MedChemExpress. Topotecan (S9321), ATR inhibitor, Berzosertib (S7102), Carboplatin (S1215), Cisplatin (S1166), Mitomycin C (S8146), and Doxorubicin (S1208) were purchased from Selleckchem.

### Organoid drug response assays

PDX tumors and organoids were dissociated into single cells, mixed with Matrigel, and plated at 1000 cells/well in 384 well poly-d-lysine coated plates (Corning). Drugs were serially diluted in organoid culture media and a total volume of 30 ul per well was added. Organoids were treated with the small molecule inhibitors, B7H3-PBD or R347-PBD (control ADC) three times for 10 days (**Figure 3A**). Thereafter, cell viability was quantified with CellTiter Glo 3D. Luminescence was measured using Infinite M200 Pro microplate reader (Tecan). Each experiment included five replicates. All treatments were repeated in two or more biologically independent experiments. Normalized AUC (nAUC) was calculated for each experiment as a ratio of AUC for B7H3-PBD and R347-PBD dose-response curves. Median nAUC and interquartile range was then calculated across independent experiments for each organoid model. Median nAUC was used for all further analysis. GraphPad Prism 9 was used to calculate IC_50_ and maximum drug effect (MaxR) which is shown as % viable cells at the highest concentration of 4ug/ml.

### Simple Western Protein quantification

LuCaP PDXs’ tissue lysates were prepared using RIPA buffer containing protease and phosphatase inhibitors. Protein samples were separated based on size using a capillary-based automated protein analysis system (Protein Simple Western Technology, Peggy Sue). Protein separation, immunodetection, and analysis steps were performed automatically by the protein analysis system. Protein bands were analyzed with the Compass software. The observed band size of B7H3 protein was variable between 120-150 kDa. GAPDH was used as the internal control. The B7H3 protein was quantified by normalizing B7H3 band intensity to the housekeeping protein, GAPDH. For the figures, the values were scaled by a factor of 10. See supplementary methods for additional details.

### Immunoblots

LuCaP PDXs’ tissue lysates were prepared as above. Organoid pellets were lysed in warm 1% SDS buffer with protease and phosphatase inhibitors. To shear the released DNA/RNA and reduce sample viscosity, 25 U/ul Benzonase nuclease (Sigma, E1014) was added per sample. The samples were incubated at 37 degrees for 30 minutes. Samples were then sonicated to shear leftover DNA using Bioruptor® Pico sonication system (Diogenode, sonication cycle: 30 sec ON/30 sec OFF, total sonication time: 5 cycles, temperature: 4 C). Protein concentration was determined by BCA assay (ThermoFisher, 23225). 10-15 ug of protein lysates were run on 4– 20% Criterion™ TGX™ Precast Midi Protein Gel (Bio-Rad) and then transferred to polyvinylidene difluoride membranes. Membranes were blocked in TBS with 0.1% Tween-20, 5% blotting grade blocker (Bio-Rad, 1706404XTU). The primary antibodies used are listed in Table S1. Clarity or Clarity Max™ Western ECL Substrate (Bio-Rad 1705061, 1705062S) was used for imaging the blots. Blots were imaged on Bio-Rad ChemiDoc Touch Imaging System.

### Flow cytometry

For B7H3 cell surface expression analysis, organoids were first dissociated into single cells using Accutase (STEMCELL Technologies, 07922). The resulting single cells were resuspended in PBS and stained with 1:1000 Zombie Violet™ dye (Biolegend, 423113) for 15 minutes to exclude dead cells. Cells were pelleted and resuspended in 100 ul staining buffer (BioLegend, 420201). Fc receptors were blocked using Human TruStain FcX™ blocking solution (BioLegend, 422301) for 5 min at room temperature. Thereafter, cells were stained with human-EpCAM-APC (Miltenyi Biotec, 130-113-260) and CD276-PE (BioLegend, 331606) for 20 min at 4°C. The stained cells were washed, fixed in 4% paraformaldehyde for 10 mins, and stored in staining buffer at 4°C up to 3-5 days. Flow cytometric analysis was done on BD LSR Fortessa cell analyzer using the BD FACSDiva software. Unstained cells and cells stained with isotype control-PE/APC antibodies were used for gating live cells and the false-positive peak, respectively. Median Fluorescent intensity(MFI) and % positive cells were analyzed using FlowJo software. % Positive cells were calculated by Overton cumulative histogram subtraction algorithm in Flowjo. Cell sorting was performed on FACSAria cell sorter (Becton Dickinson) using FACSDiva software.

### CRISPR/Cas9 experiments

We used three organoid models for CRISPR experiments: LuCaP 145.2, LuCaP 173.1, and NCI-PC155. Two vector system (lentiCas9-Blast, lentiGuide-Puro) was used. First, Cas9 expressing organoid lines were generated by lentiviral transduction of Cas9 plasmid (lentiCas9-Blast, addgene; 52962) and selection with Blasticidin for 10-14 days. Individual sgRNAs were cloned into lentiGuide-Puro plasmid (NCI core facility). Cas9 expressing organoids were then transduced with lentiviral sgRNA (sgB7H3 #1; CAACCGCACGGCCCTCTTCCCGG) and selected with 1 ug/ml puromycin for 3 days. All lentiviral transductions were done by spin infection at 1000 x g, for 90 mins at 32°C.

Since clonal culture was not possible from organoids, B7H3 sgRNA transduced LuCaP 145.2 organoids were processed into single cells and FACS sorted to enrich for B7H3 knockout cells. For later experiments, FACS sorted B7H3-WT and B7H3-KO cells were maintained in 3D organoid cultures. B7H3 knockdown was periodically confirmed by western blotting.

Details of the CRISPR/Cas9 dropout screen are provided in supplementary methods.

### Preclinical trials

Male NSG mice were subcutaneously injected with 2 million processed LuCaP PDX tumor cells or implanted with tumor pieces. When tumor exceeded ∼150 mm3, mice were randomized into three groups - B7H3-PBD, R347-PBD, and vehicle. 1mg/kg of the B7H3-PBD or R347-PBD was administered intraperitoneally weekly, for two weeks. Tumor growth (mm^3^) and body weight were measured twice a week. Tumor growth is shown for the individual mouse from the time of first treatment. Mice were euthanized if any of the two tumor diameters reached 2cm or if their health was compromised.

For establishing LuCaP 136 metastasis model, processed PDX cells were first transduced with a luciferase reporter plasmid and re-injected subcutaneously in NSG mouse. The resulting tumor was processed, and 100,000 single cells were injected intracardiacally. Tumor growth was measured by bioluminescence imaging. Treatment was started when the luminescence signal reached ∼3 x 10^7^.

### Organoid derived xenografts

Organoid derived xenografts (ODXs) were established in NOD scid gamma (NSG) mice as previously described (21). The subsequent subcutaneous tumor was collected, processed and 1-2 million cells, mixed with Matrigel in 1:1 ratio, were injected subcutaneously per mice for the experiment. Mice were randomized into three treatment groups (B7H3-PBD, R347-PBD, and vehicle) when the tumors reached an average volume of 150-250 mm^3^. The treatment was started as described above.

### Toxicology studies with B7H3-PBD-ADC

In addition to evaluating changes in body weight after administration of the ADC, we also performed full necropsy on day 8 and day 30 post-treatment to evaluate acute and delayed toxicity, respectively. Tumor bearing mice (LuCaP 136 PDX model) were administered two intraperitoneal doses of 1mg/kg B7H3-PBD-ADC or control R347-PBD-ADC, given weekly for two weeks. A gross necropsy was performed and a standard list of organs, including brain, lung, liver, kidney, spleen, thymus, testes, heart, and bone, were embedded in paraffin, sectioned, stained with hematoxylin and eosin, and examined microscopically by a board-certified veterinary pathologist. Additionally, to assess accumulation of ADC in different organs, IHC for human IgG was performed on sections of liver, adrenal, kidney, and prostate collected on day 8 or day 30 post-treatment from all animals. IHC assay positive control tumor tissue displayed appropriate IgG membrane immunoreactivity and no signal was observed in negative control tissues.

### RNA-sequencing

RNA-seq was performed on the mPC organoid models on Illumina NovaSeq_S2 Flowcells using rRNA-depleted RNA and paired-end reads. Data were preprocessed and filtered using our previously described pipeline (21). RNA sequence reads were aligned with STAR to the human genome, hg19. The featureCounts program from the subread package was applied to compute the raw reads count. All further analysis was performed in R. Data were TMM normalized to convert the raw read count to log_2_ transformed CPM (counts per million) using EdgeR. Median gene expression values were used for the biological replicates. Normalized log_2_CPM values were used for all subsequent analyses.

ConsensusDE package was used to perform differential expression analysis with three different algorithms (EdgeR, DESeq, and Voom) simultaneously. Genes were considered differentially expressed if adjusted Benjamin-Hochberg false discovery rate (FDR) was less than 0.05 for all three methods.

### Gene signature scores

Single sample gene set signature scores were calculated using GSVA with default parameters (34), using all MSigDB gene sets and interferon signature score (IFN score) refined for prostate cancer. Gene signature scores (IFN score, Replication stress score, AR score, and RB score) used in this study are described in detail in the supplementary methods.

### Clinical data analyses

SU2C/PCF 2019 data was downloaded from cbioportal (3). Log_2_(FPKM+1) mean centered data was used. Since PolyA and capture sequenced RNAseq data are highly concordant (28), we merged the two datasets to have unique 328 samples. Samples with at least 30% tumor content (n = 303/328) were kept for analysis. Cbioportal data was merged with curations of the *TP53/RB1* status and AR/NE state from Nyquist et al 2020 (28). IFN score and RB1 score were calculated using GSVA with method argument as “ssgsea” (34). For analysis of *SLFN11* status, data was normalized using ordered quantile normalization with the orderNorm function in R.

### Statistics

R package, ggpubr, and GraphPad Prism were used for all analyses. Pearson’s correlation coefficient was calculated for scatterplots using ggscatter function in R. The nonparametric Kruskal Wallis test was used to compare multiple groups. For pairwise comparison between groups, Wilcoxon test was used with *P* value adjusted for multiple testing using the Holm method. Differences were considered significant for two-tailed *P* values of < 0.05. For correlation analysis, Benjamini and Hochberg method was applied for multiple hypothesis test correction with adjusted *P* value (FDR) ≤ 0.05 indicating statistical significance.

For in vivo metastasis experiment, treatment response was compared using GraphPad Prism by two-way Anova. Kaplan-Meier survival analysis was performed, and the survival rate was compared by Log-rank test. Data are expressed as the mean ± SEM, with *P* < 0.05 considered statistically significant.

Heatmap was generated by using the R package ComplexHeatmap.

### Study Approval

Propagation of LuCaP PDX tumors and in vivo drug studies were performed at the NCI under NCI Animal Care and Use Committee approved protocol. Tumors were periodically validated using STR analysis by Laragen Inc. All animal procedures were approved by NCI Animal Care and Use Committee. For organoids (NCI-PC44 and NCI-PC155) derived from patient biopsies, patients provided informed consent, and samples were obtained from the NIH clinical center under NIH Institutional Review Board approval in accordance with U.S. Common Rule. All rapid autopsy tissues were collected from patients who had signed written informed consent under the aegis of the Prostate Cancer Donor Program at the University of Washington (2).

## Authors contributions

Conceptualization: SA, EMH, KK

Methodology: SA, LF, KM, JY, PSN, AM, CH, EMH, KK

Investigation: SA, LF, KM, JY, ANA, EC, MR, LT, RD, IC, JL, MB, PSN, ISS, AM, CH

Visualization: SA, JB, ATK, BJC, AGS, ISS, PSN, AM, CH

Funding acquisition: EMH, KK Supervision: EMH, KK

Writing – original draft: SA, KK

Writing – review & editing: SA, EMH, PSN, KK

## Acknowledgments

The authors wish to express their gratitude to the patients and the families of the patients who contributed to this study. We thank Yves Pommier for reviewing the manuscript. We thank Alison Smith and Kelly Ryan in the Tumor Targeted Delivery Chemistry group at AstraZeneca. We also thank all the NCI/CCR core facilities including LGCP microscopy core, Flow cytometry core for assistance with cell sorting, Genomics core for performing RNA sequencing, Genome Modification core for designing CRISPR guides, and Nanoscale Protein Analysis Section of the CCR Collaborative Protein Technology Resource for performing Simple Western assays.

## Data and materials availability

All data are available in the main text or the supplementary materials.

List of Supplementary Materials

Materials and Methods

Details of anti-B7H3 ADC generation Simple Western protein quantification details IF staining

Immunohistochemistry CRISPR/Cas9 dropout screen Gene signature scores

Figure S1. B7H3 expression is mPC patient samples, PDXs, and organoids.

Figure S2. B7H3 FACS analysis and correlation of B7H3 expression and AR score. Figure S3. B7H3-PBD response in organoid models and B7H3 knockout assays.

Figure S4. Biomarker analysis of B7H3-PBD response.

Figure S5. Assay for ATR activity in selected organoids following treatment with chemotherapeutics.

Figure S6. In vitro and in vivo safety profile of the B7H3-PBD-ADC.

Figure S7. Analysis of SU2C data to predict likely responders based on identified biomarkers. Table S1. List of antibodies.

Data file S1. Descriptive data for B7H3-PBD response in 26 organoid models.

Data file S2. Differentially expressed genes between responder and Non-responder ARPC models.

References (*34–36*) only cited in the supplementary materials.

