## Supplementary material for "Tumor-derived biomarkers beyond antigen expression enhance efficacy of CD276/B7H3 antibody-drug conjugate in metastatic prostate cancer": B7H3_supplementary_data

**Supplementary methods**

**Details of anti-B7H3 ADC generation**

B7H3 antibodies were isolated from naive human phage display libraries using phage display soluble selections, performed according to standard methods (Vaughan 1996; Lloyd 2009). Three rounds of selection were performed against recombinant human B7H3 4Ig_avitag_10His protein, immobilised on wells of a MaxiSorp® microtitre plate (Nunc) overnight at 4°C. Round 2 and round 3 selection outputs were evaluated for diversity, sequence uniques, and specificity, using phage ELISA to assess binding specificity across biotinylated human 4Ig, human 2Ig, and cyno 4Ig proteins.

Forty-four sequence-diverse clones with robust B7H3 binding were selected from round 2 outputs, which had the highest sequence diversity. His-purified single chain variable fragments (scFv) were then screened for binding to a range of B7H3-expressing tumor cell lines. From these binding data, 20 were prioritized for conversion to HuIgG1 and site-specific cysteine conjugation to a PBD payload at DAR2, with subsequent screening for in vitro ADC cytotoxicity activity across 20 tumor cell lines.

Two molecules with the best cytotoxicity profile were chosen for affinity maturation, again using phage display soluble selections and targeted CDR3 mutagenesis. ScFv fragments were screened for improvements in Hu 4Ig biochemical binding, with sequence uniques profiled on 4Ig, 2Ig and species variants of B7H3 protein. Subsequent conversion to HuIgG1 was followed by further protein & cell binding studies, with conjugation and cytotoxicity for the most promising molecules.

### Simple Western protein quantification details

Simple Western analysis was performed on Peggy Sue instrument according to the ProteinSimple user manual. Cell lysates adjusted to contain the same amount of protein were mixed with sodium dodecyl sulfate master mix (containing dithiothreitol (DTT) and fluorescent molecular weight markers, ProteinSimple), and were heated at 70°C for 10 min before loading to the instrument for fully automated analysis. Proteins (40 ng loaded) were separated based on molecular weight while migrating through the separation matrix; the separated proteins were immobilized to the capillary wall using UV light, and incubated with blocking reagent (ProteinSimple), followed by immunoprobing with respective primary antibodies (CD276 1:50 (AF1027, R&D) and GAPDH 1:100 (MAB374, Millipore) and HRP-conjugated anti-goat (1:100) or anti-mouse (1:600) secondary antibodies (Jackson ImmunoResearch). A 1:1 mixture of luminol and peroxide (ProteinSimple) was added to generate chemiluminescence, which was captured by CCD camera. The digital image was analyzed by Compass software (vs. 5.0.1; ProteinSimple). Target protein quantities were determined by calculating the area under the peak identified as CD276 (B7H3) and GAPDH, a housekeeping protein used as loading control.

### IF staining

Organoids were dissociated and plated in 200 ul culture media (+/- doxycycline) with 2% Matrigel on poly-d-lysine coated chamber slides (Ibidi, 81201). After 24 hours, cells were fixed in 4% paraformaldehyde for 10 mins and stained with RB1 antibody and DAPI. Images were taken with Zeiss Axioscan.Z1 slide scanner.

### Immunohistochemistry

For tissue microarray (TMA) slides automated IHC was performed on the VENTANA Discovery Ultra (Ventana Medical Systems Inc) autostainer. Onboard deparaffinization was conducted in DISCOVERY Wash buffer (VMSI, 950-510). Subsequent heat-induced epitope retrieval (HIER) was performed in DISCOVERY CC1 solution (VMSI, 950-500). Tissue microarray sections were incubated with B7H3 recombinant rabbit monoclonal antibody (Sigma Aldrich, SP206) at a dilution of 1:500 in Ventana Antibody Diluent with Casein (VMSI, 760-219). The primary antibody was bound with DISCOVERY anti-Rabbit HQ secondary antibody (Ventana), followed by DISCOVERY Anti-HQ HRP (VMSI, 760-4820) enzyme conjugate. The antibody complex was visualized using the ChromoMap DAB detection Kit (VMSI, Cat# 760-159). Hematoxylin II (VMSI, 790-2208) and Bluing Reagent (VMSI, 760-2037) were used to counterstain the sections.

**CRISPR/Cas9 dropout screen**

For the mini CRISPR/Cas9 dropout screen, Cas9 expressing organoids were dissociated and transduced with a custom lentiviral library of 16 sgRNAs which included two sgRNAs targeting B7H3 (sgB7H3 #1; CAACCGCACGGCCCTCTTCCCGG, sgB7H3 #2; CTCAGGGTAGCCCCGGTAGCTGG) along with one positive control sgRNA (U2AF1; GTCATGGAGACAGGTGCTCT) and two non-targeting control sgRNAs (ACTGCTCCCGGTCGCCCCTC , CGCACGACCATTGCTGCTGC). Transduction was done at a low MOI of 0.3-0.5 to maximize the likelihood of integrating one sgRNA per cell. Transduced cells were plated in 3D in 5% Matrigel on ultra-low attachment plates (Corning). After 24 hours, organoids were selected with 1ug/ml puromycin for 3 days.

Organoids were then passaged, and cell pellets were collected at the indicated time points up to 22 days. Day 0 indicates the first time point after puromycin selection. A small library of 16 sgRNAs allowed >10,000x library coverage, with 200,000 cells pelleted at each time point for harvesting gDNA. All experiments were done in duplicate.

gDNA extraction and subsequent PCR amplification and purification of the sgRNA bar-coded regions were done according to the BROAD's protocol (https://portals.broadinstitute.org/gpp/public/resources/protocols). PCR libraries were sequenced on an Illumina MiSeq 2x150 bp (Genewiz). The sequencing data was analyzed using the MAGeCK package. Fold change of sgRNA read counts was calculated between the samples and the baseline Day 0 samples. The results shown are for two B7H3 sgRNAs in comparison with the positive control sgRNA and two negative control sgRNAs.

**Gene signature scores**

**(A) IFN signature score**

Normalized $log_2$CPM gene expression values were converted to modified Z score (ZMAD) to achieve normal distribution. We then evaluated the expression of 49 interferon (IFN) genes from the previously published IFN-related DNA damage resistance signature (IRDS) for breast cancer (35). As a first pass for deriving a prostate specific IFN signature, we focused on genes that were detected in all LuCaP samples and calculated the IFN score. Next, we evaluated whether any of the other interferon genes, excluded in breast cancer signature, are correlated with IFN score in our cohort. In this case, we selected IFN genes that had adjusted p value $\leq$ 0.05, a correlation coefficient greater than 0.5, and were a member of the Hallmark IFN alpha or Hallmark IFN gamma gene signature. A list of significantly correlated genes was defined as prostate cancer specific IFN signature and prostate specific IFN score was calculated. We further refined the IFN signature by again performing correlation analysis with prostate specific IFN score, but this time, using all genes. Additionally, we tightened our filtering criteria by using adjusted p value $\leq 0.05$ and lower bound of the confidence interval $> 0.5$ for filtering as opposed to the correlation coefficient. This resulted in 47 gene signature which was used to calculate the refined IFN score referred as "IFN Score" in the text and figures.

**(B) Replication stress score**

Replication stress signature score was generated using methodology previously described (36). Briefly, we selected Reactome gene sets for signatures related to DNA repair, replication, and cell cycle to define biological processes indicative of replication stress response. Initially, organoid RNAseq data was filtered by genes involved in the above selected pathways. Weights for the genes were generated by implementing principal component analysis across all models and derived from the first principal component. These weights were then applied across all models via a dot product and summated to create replication stress (RepStress) scores for each model.

**(C) AR score and RB score**

AR score was calculated as described previously (37). RB.CRPC genes from McNair et al (38) were used as a gene set to calculate RB signature score using the GSVA function with default parameters from the GSVA R package (34).

### Supplementary figures

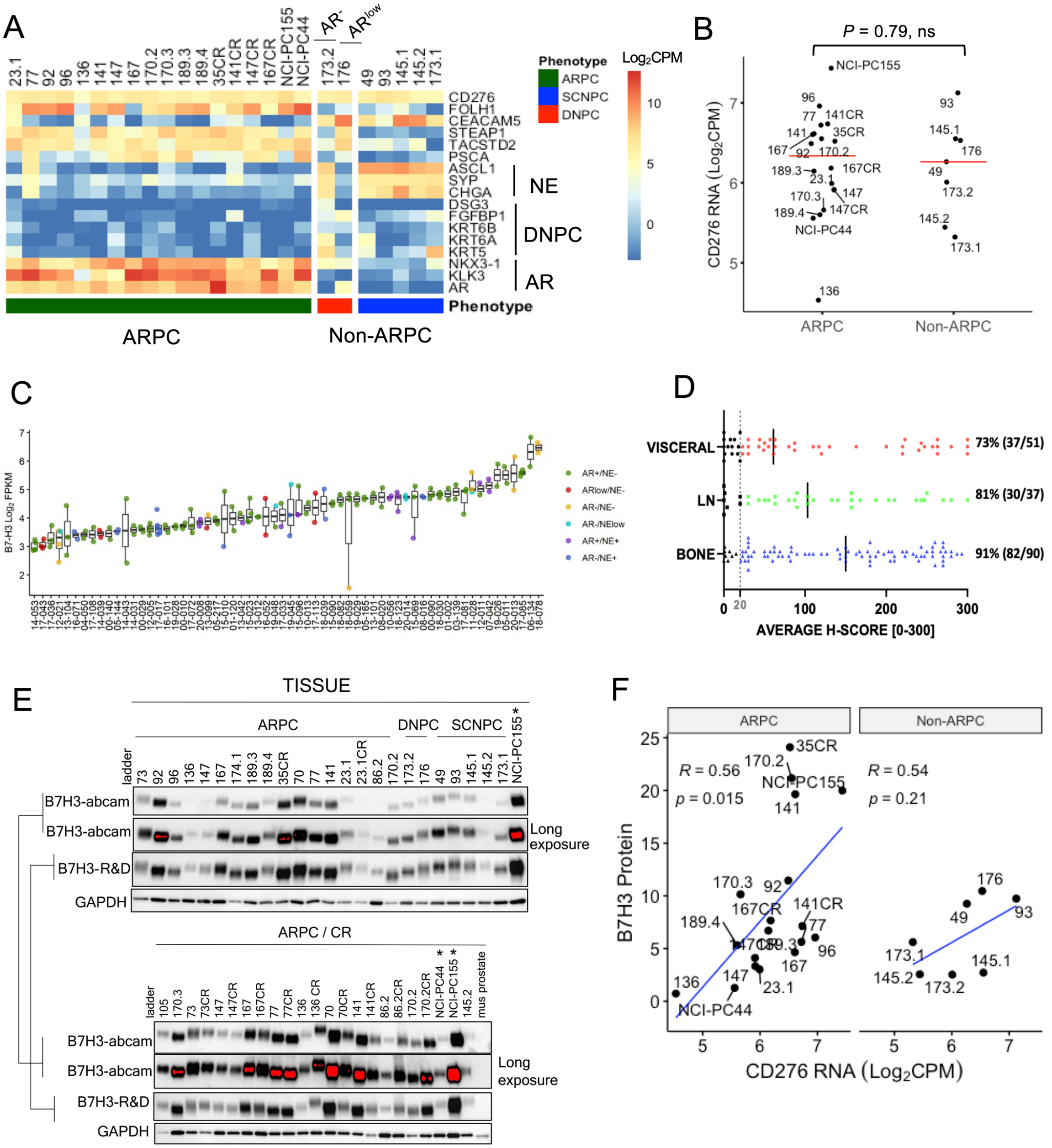


Sup Fig 1

**Figure S1. CD276/B7H3 expression is mPC patient samples, PDXs, and organoids.** (Related

**to Figure 1). (A)** Molecular profile of organoid models tested in this study based on AR-, DNPC-, and neuroendocrine (NE) associated genes. *CD276/B7H3* and *FOLH1/PSMA* transcript levels are shown as $Log_2CPM$. ARPC = AR-active adenocarcinoma (Intact and experimentally castrate resistant), Non-ARPC = SCNPC (LuCaPs 49, 93, 145.1, 145.2, 173.1) and $AR^{NEG/LOW}$ $NE^{NEG}$ (LuCaPs 173.2 and 176, respectively, grouped as DNPC) (2) phenotypes. Transcript data not available for 23.1CR and 77CR organoids. **(B)** Dot plot for B7H3 RNA expression in mPC organoids based on phenotypes. The red line indicates median B7H3 expression. **(C**) Intra-individual inter-tumor variation in CD276/B7H3 expression in patients with metastatic prostate cancer. *CD276/B7H3* transcript abundance determined by RNA sequencing analysis of metastatic prostate tumors. Transcript levels are shown as Log2 FPKM. Boxplots include patients with at least two tumors profiled (149 tumors from 62 patients) and are ordered by per-patient median log2 FPKM gene expression**.** (**D**) Distribution of B7H3 protein expression in metastatic tumors from different sites. Dashed line shows H-score of 20. Solid line shows the mean H-score. **(E)** Western blot analysis of PDX samples (n=35), patient derived organoids (n =2, indicated by *) and mouse prostate tumor (negative control). Similar B7H3 expression pattern is shown with two different B7H3 antibodies, Abcam (ab134161) and R&D (AF1027). GAPDH was used as a loading control. Oversaturated protein bands from long exposure are marked in red. CR = experimentally castrate resistant (**F)** Pearson correlation between B7H3 protein and RNA in the ARPC and non-ARPC models (ARPC models: r = 0.56, n =18, *P* = 0.015; Non-ARPC models: r = 0.54, n =7, *P* = 0.21).

136

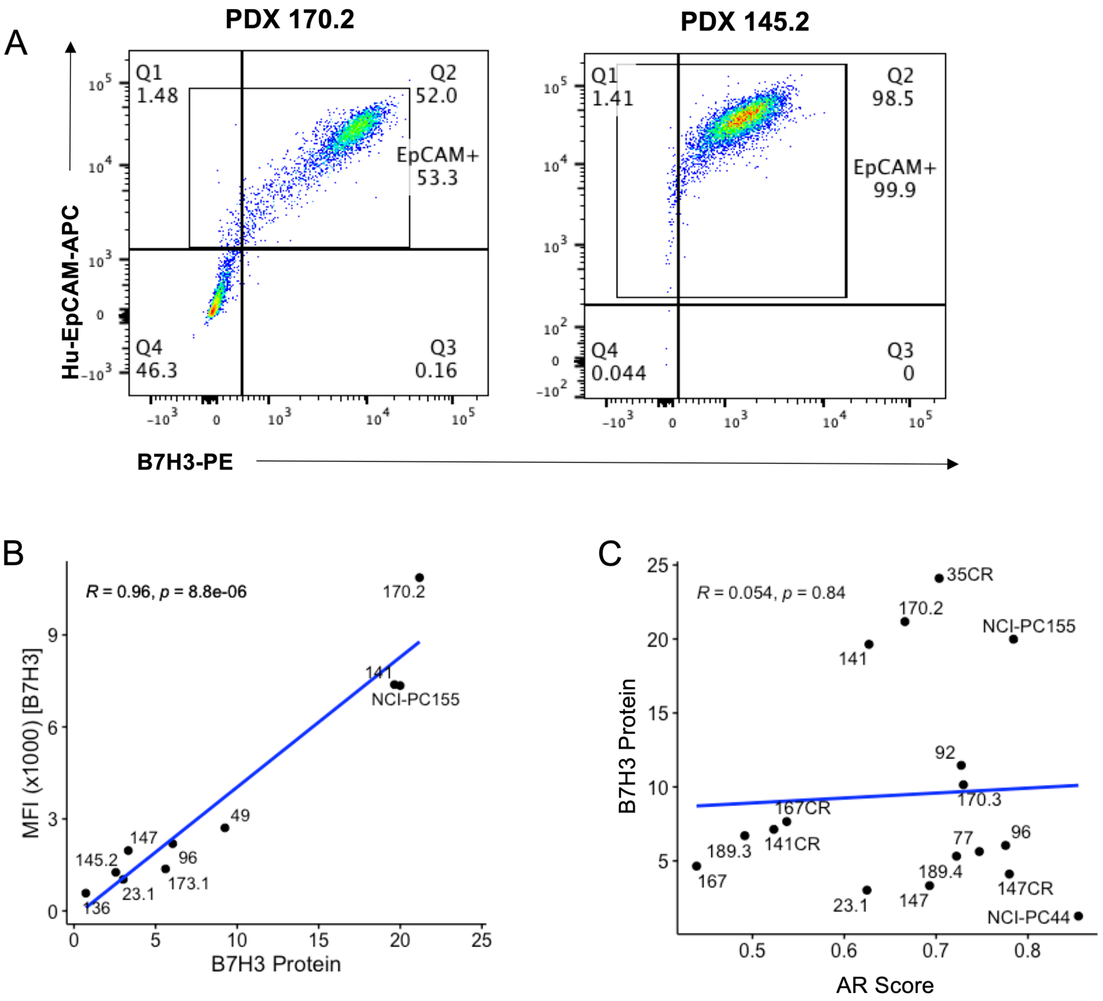


Sup Fig 2

137

**Figure S2. B7H3 FACS analysis and correlation of B7H3 expression and AR score (related to Figure 2). (A)** Representative figures showing FACS gating strategy for EpCAM+/ B7H3+ cell population in two organoid models, PDX 170.2 ARPC (left) and PDX 145.2 non-ARPC (right). **(B)** Scatter plot for Pearson correlation between B7H3 cell surface expression by flow cytometry (MFI) and total protein expression by Simple Western. **(C)** Plot of AR score and B7H3 protein for PDX models. $P = 0.84$, not significant.

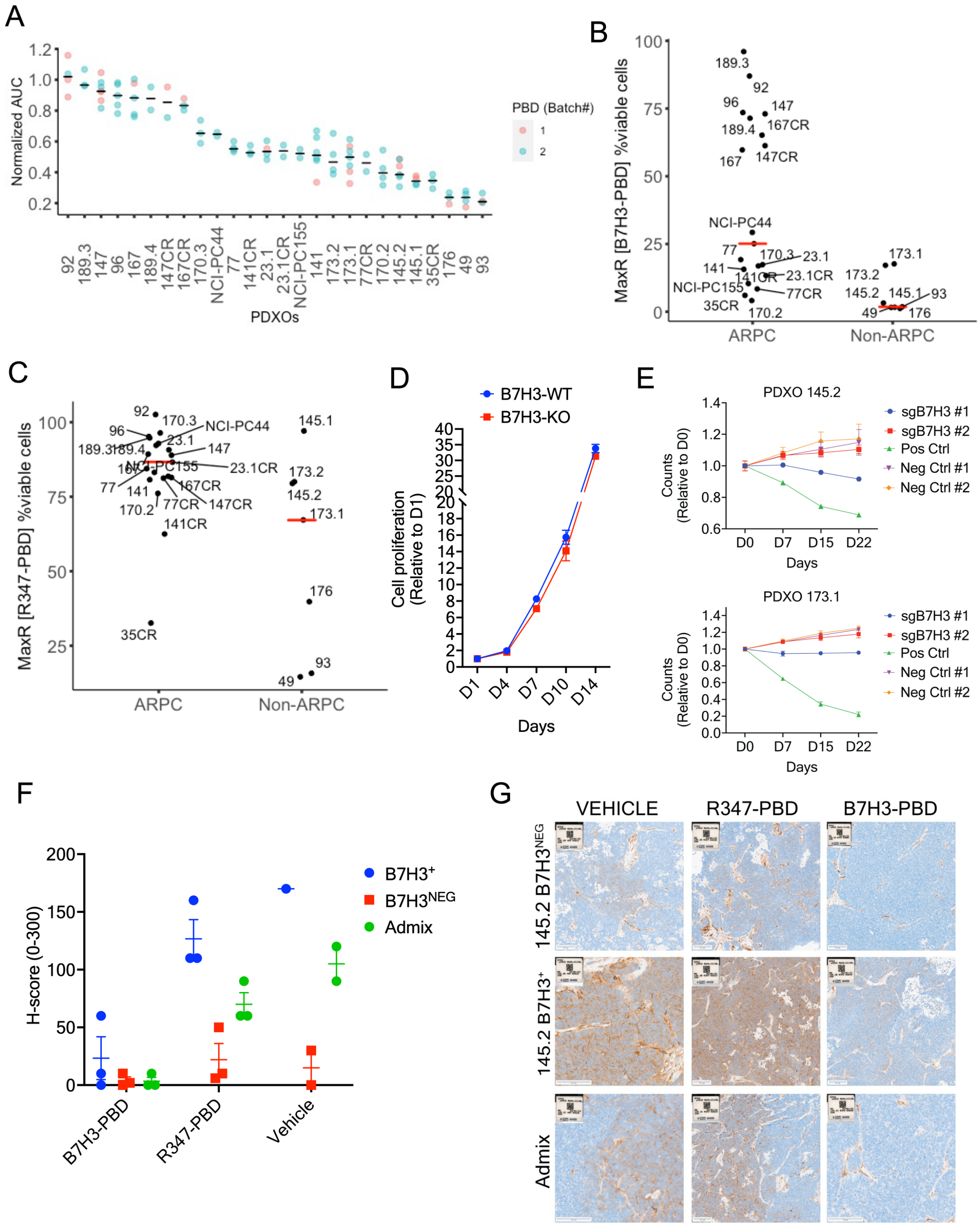


Sup Fig 3

**Figure S3. B7H3-PBD response in organoid models and B7H3 knockout assays (related to**

**Figure 3). (A)** Normalized AUC for each organoid model is plotted. Dots indicate number of independent biological replicates tested for each model with the two different batches of ADC (Batch #1 and #2). **(B-C)** For each organoid model, median value of biological replicates is plotted to show maximum response (MaxR) at 4 ug/ml dose of **(B)** B7H3-PBD-ADC and **(C)** R347-PBD-ADC . MaxR is shown as % viable cells left at the tested maximum concentration of 4ug/ml. **(D)** Growth comparison of B7H3-WT(B7H3$^{+}$) and B7H3-KO(B7H3$^{NEG}$) LuCaP 145.2 organoids at indicated time points by 3D CellTiter Glo; n =5 replicates for each. Error bars indicate the SEM. **(E)** CRISPR dropout screen in LuCaP 145.2 and LuCaP 173.1 organoids. Depletion of two B7H3 guides (sgB7H3 #1, sgB7H3 #2), two negative control guides, and a positive control single-guide RNA is shown at indicated time points relative to day 0 (D0).**(F-G)** Immunohistochemical assessments of B7H3 protein expression in xenografts from sorted B7H3$^{+}$, B7H3$^{NEG}$, and admix (mix of B7H3$^{+}$ and B7H3$^{NEG}$) LuCaP 145.2 cells treated with vehicle, B7H3-PBD-ADC or R347-PBD-ADC. All tumors were collected when mice in vehicle cohort reached endpoint. n=3 per group, except vehicle treated group (B7H3$^{+}$, n=1; B7H3$^{NEG}$, n =2; admix, n=2) **(F)** Dot plot shows H-score. Solid line shows the mean H-score. (**G)** Representative IHC images are shown.

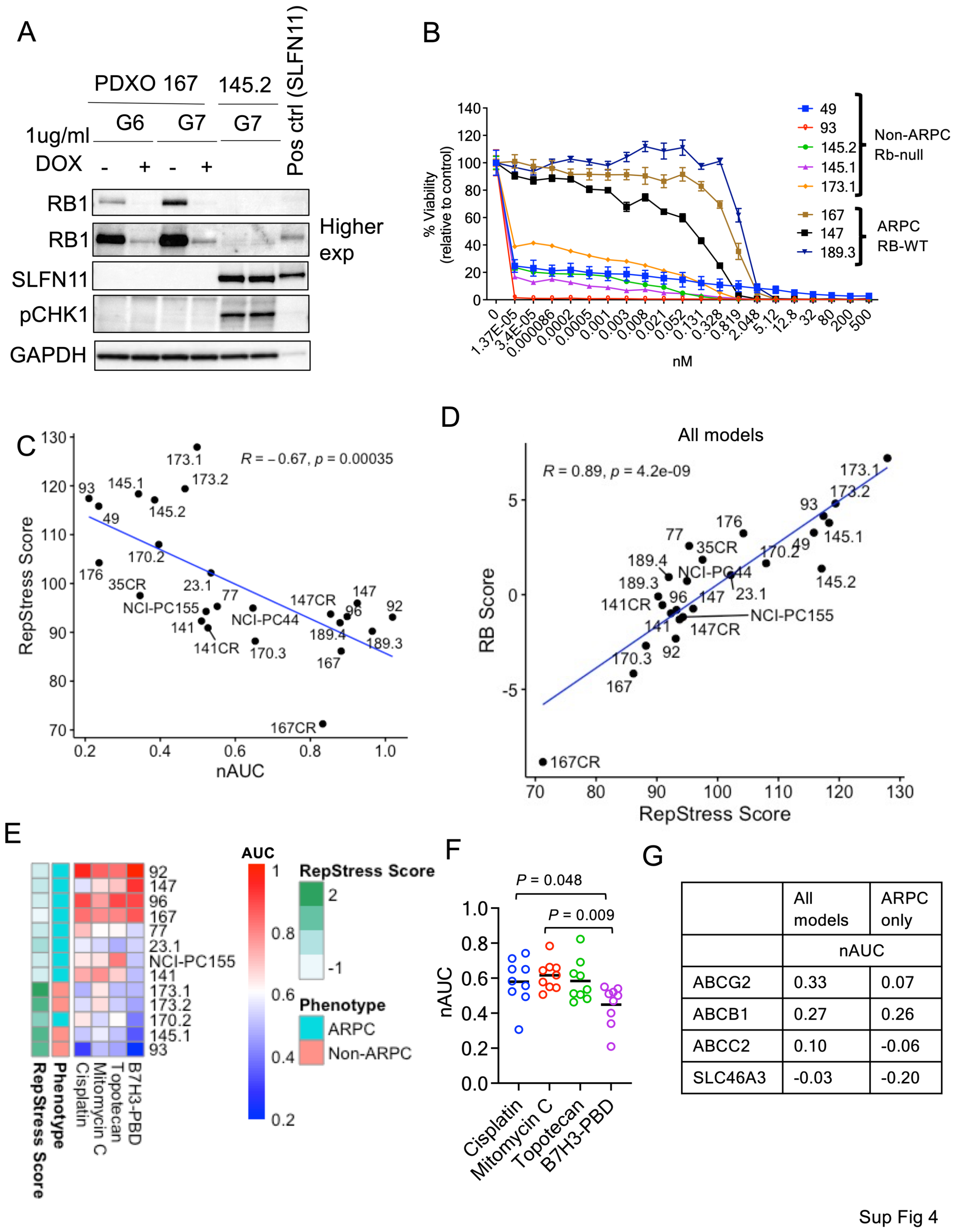


| | All models | ARPC only |
|---|---|---|
| | nAUC | |
| ABCG2 | 0.33 | 0.07 |
| ABCB1 | 0.27 | 0.26 |
| ABCC2 | 0.10 | -0.06 |
| SLC46A3 | -0.03 | -0.20 |

**Figure S4. Biomarker analysis of B7H3-PBD-ADC response. (Related to Figure 4 and 5)**

**(A)** Immunoblot confirming RB1 knockdown with doxycycline inducible shRNA in PDX 167 organoid model. PDX 145.2 organoids ($RB1^{loss}$) is used as a RB1 negative control. Effect on SLFN11 and pCHK1 (Ser345) expression is also shown. SLFN11 positive control lysate is from 293T cells transfected with SLFN11 expression vector. GAPDH is used as a loading control. **(B)** Dose response curve for free PBD dimer comparing sensitivity of RB1-wt and RB1-null models. **(C)** Plot comparing B7H3-PBD-ADC nAUC and RepStress score. Pearson's correlation coefficient r = -0.67, *P* = 0.00035. **(D)** Pearson correlation between RB1 score and RepStress score across all models. r = 0.89, *P* = 4.2e-09. **(E-F)** Heatmap (**E**) and dot plot (**F**) of normalized AUC (nAUC) values for organoid models in response to drugs targeting replication stress; B7H3-PBD, topotecan, cisplatin, and mitomycin C. Black line in the dot plot (**F**) indicates mean nAUC value. **(G)** Pearson's correlation analyses between RNA levels of previously identified biomarkers and PBD sensitivity. r values are shown. *P* = not significant.

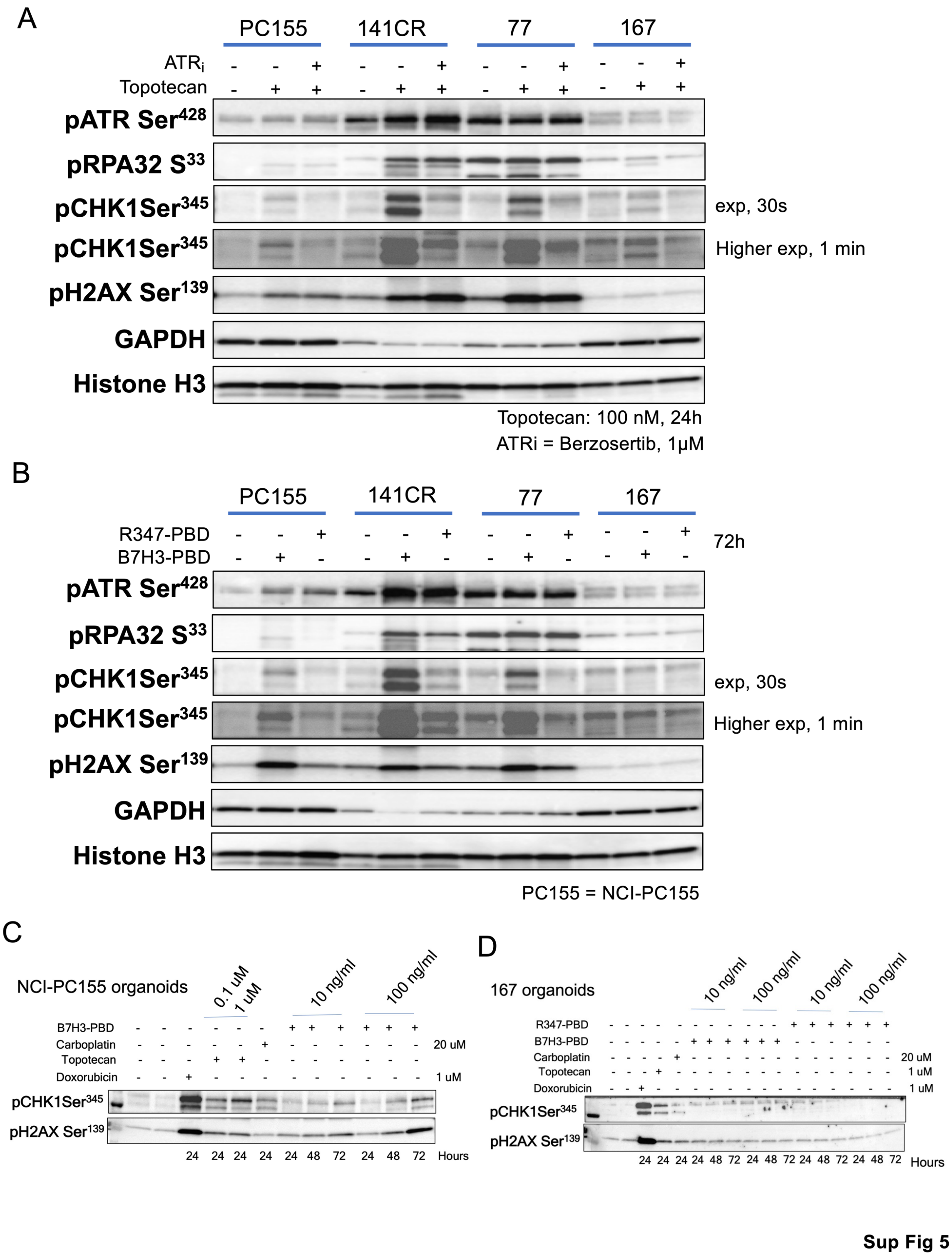


**Figure S5. Assay for ATR activity in selected organoids following treatment with**

**chemotherapeutics. A and B:** ATR activity was measured by assessing pATR-$Ser^{428}$, pRPA32-$S^{33}$ and pCHK1-$Ser^{345}$ levels. DNA damage in response to the treatment was measured by pH2AX $Ser^{139}$ levels. GAPDH and total histone H3 were used as loading controls. **(A)** Organoids were treated with topotecan (100 nM) for 24 hours in the presence or absence of $ATR_i$ (Berzosertib). **(B)** Organoids were treated with B7H3-PBD-ADC or R347-PBD-ADC for 72 hours. **(C-D)** Expression of pCHK1-$Ser^{345}$ and pH2AX $Ser^{139}$ in (C) NCI-PC155 organoid model and (D) 167 organoid model in response to treatment with B7H3-PBD-ADC, R347-PBD-ADC, carboplatin, topotecan, and doxorubicin. Concentration and length of treatment is indicated for each drug.

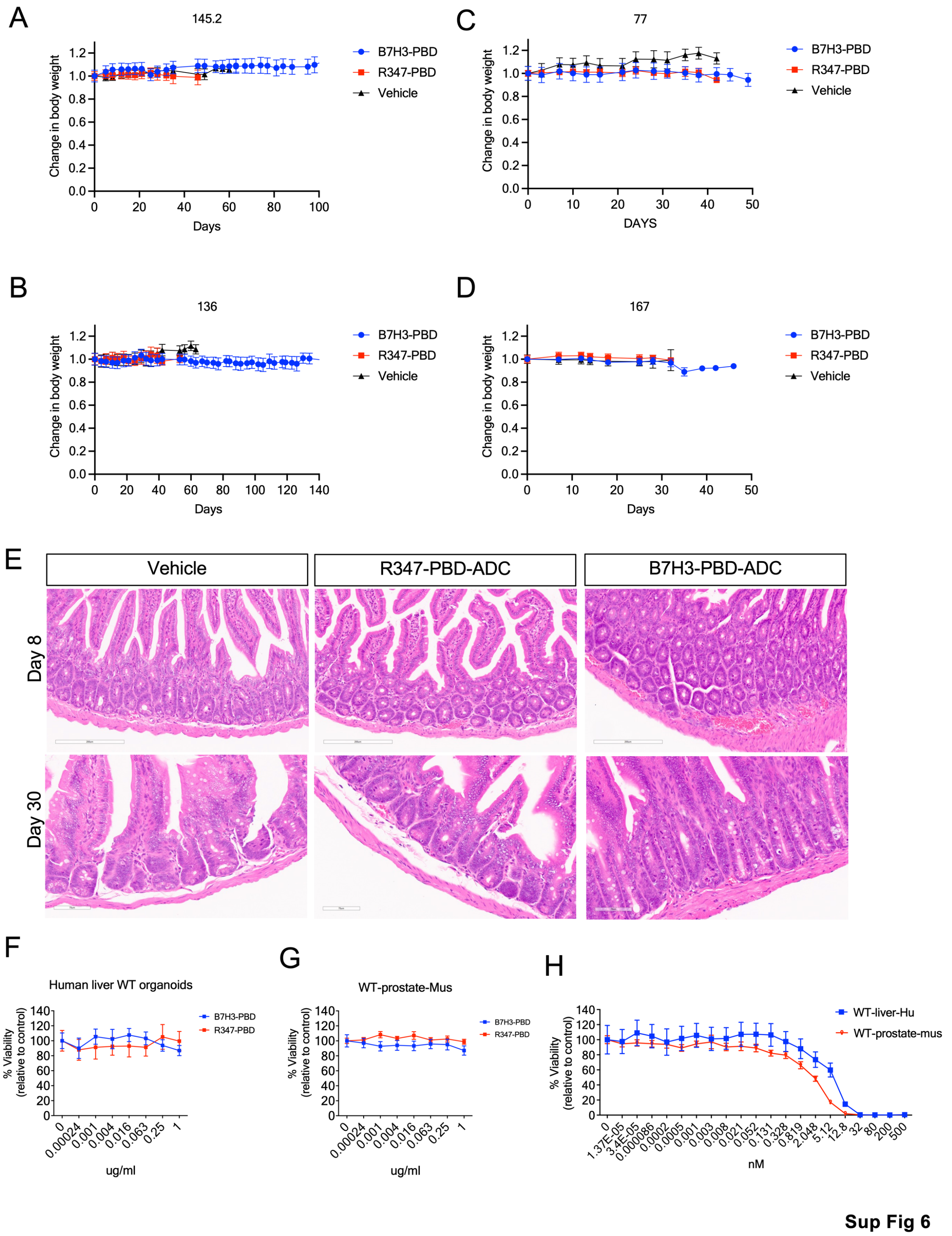


**Figure S6. In vitro and In vivo safety profile of ADC. (A-D)** Body weight for mice treated with B7H3-PBD-ADC, R347-PBD-ADC or vehicle (Related to figure 5). **(E)** H&E sections of small intestine collected on day 8 and day 30 post-treatment. **(F-H)** in vitro toxicity analysis of B7H3-PBD-ADC, R347-PBD-ADC, and free PBD dimer. (**F**) Normal human liver organoids and (**G**) wild type mouse prostate organoids were treated with the ADC for 10 days. Dose response curves are shown. (**H**) Dose response curve for free PBD dimer in normal human liver and normal mouse prostate organoids.

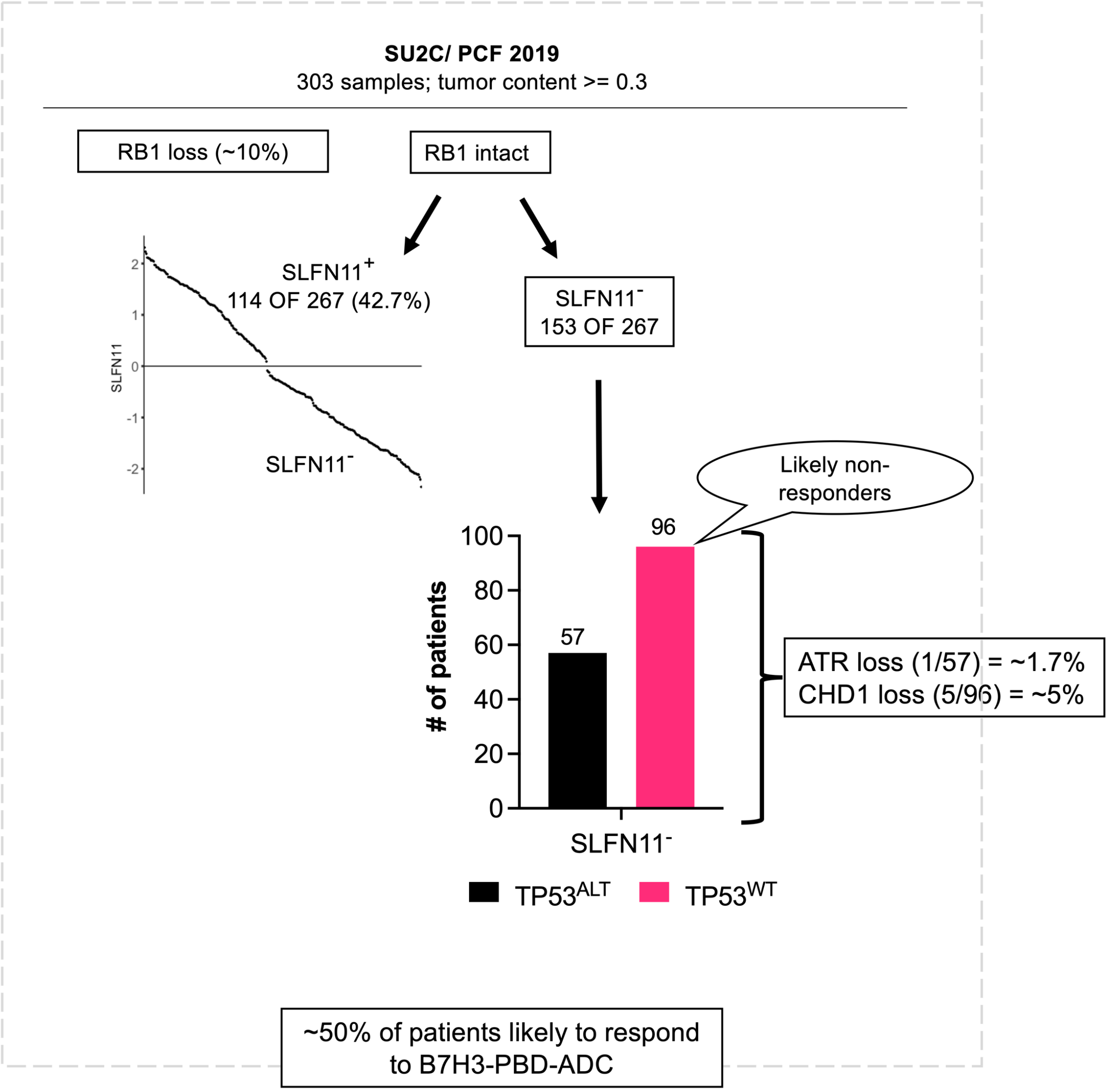


**Figure S7. (A)** Analysis of SU2C clinical data to predict likely responders based on identified biomarkers of B7H3-PBD-ADC sensitivity (**Related to figure 4H**). $Log_2$(FPKM1+1) mean centered values are plotted for *SLFN11* expression after ordered quantile normalization. Distribution of SLFN11$^-$ patients based on *TP53* genomic status is shown. TP53$^{WT}$ = at least one WT allele and absence of gain of function mutation.

**Table S1. List of antibodies used.**

| Target | Company | Catalog number | Assay |
|---|---|---|---|
| AR | Abcam | ab133273 | Western |
| p53 | Cell Signaling | 48818 | Western |
| PTEN | Cell Signaling | 9188 | Western |
| RB | Cell Signaling | 9309 | IF/Western |
| B7H3 | Abcam | ab134161 | Western |
| B7H3 | R&D | AF1027 | Western / Simple Western |
| GAPDH | Abcam | ab8245 | Western |
| SLFN11 | Cell Signaling | 34858 | Western |
| ATR | Cell Signaling | 2790 | Western |
| pCHK1 (Ser345) | Cell Signaling | 2348 | Western |
| ATM | Cell Signaling | 2873 | Western |
| pH2AX (Ser129) | Cell Signaling | 9718 | Western |
| p21 | Cell Signaling | 2947 | Western |
| CHD1 | Cell Signaling | 4351 | Western |
| KLK3 | Cell Signaling | 5365 | Western |
| pATR (Ser428) | Cell Signaling | 2853 | Western |
| pRPA32 (S33) | Bethyl labs | A300-246A | Western |
| Histone H3 | Cell Signaling | 4499 | Western |
| CD276-PE | Biolegend | 331606 | FACS |
| EPCAM-APC | Miltenyi | 130-113-260 | FACS |
